# Structure of the Human ATAD2 AAA+ Histone Chaperone Reveals Mechanism of Regulation and Inter-subunit Communication

**DOI:** 10.1101/2022.10.10.511654

**Authors:** Carol Cho, Ji-Joon Song

**Affiliations:** Department of Biological Sciences, Basic Science 4.0 Institute, KAIST Stem Cell Center, Korea Advanced Institute of Science and Technology (KAIST), Daejeon, 34141, Korea

## Abstract

ATAD2 is a non-canonical ATP-dependent histone chaperone and a major cancer. Despite widespread efforts to design drugs targeting the ATAD2 bromodomain, little is known about the overall structural organization and AAA+ domains of ATAD2. Here, we present the 3.1 Å cryo-EM structure of human ATAD2 in the ATP state, showing a shallow hexameric spiral that binds a peptide substrate at the central pore. The spiral conformation is locked by an N-terminal linker domain (LD) that wedges between the seam subunits, thus limiting ATP-dependent symmetry breaking of the AAA+ ring. In contrast, a structure of the ATAD2-histone H3H4 complex shows the LD undocked from the seam, suggesting that H3H4 binding unlocks the AAA+ spiral by allosterically releasing the LD. These findings, together with the discovery of an inter-subunit signaling mechanism, reveal a unique regulatory mechanism for ATAD2 and lay the foundation for developing new ATAD2 inhibitors.

## INTRODUCTION

The oncogene ATAD2 (ATPase family AAA+ domain-containing 2) is systematically up-regulated in various cancers of unrelated origins^1,2^, and is linked to many oncogenic transcription factors such as cMyc and E2F^3^. ATAD2 expression levels correlate with poor patient prognosis in colorectoral^4^, lung^1^, and breast cancer^5^, and knockdown of ATAD2 attenuates tumor proliferation and invasiveness^6^. Consequently, ATAD2 has been widely explored as a therapeutic target for various cancers, resulting in several potent small molecule ATAD2 inhibitors published to date^7,8^ with potentially more under development. Despite such interest in discovering ATAD2-targeted drugs, surprisingly little is known about the cellular roles of ATAD2 and the mechanism of tumor malignancy.

Although ATAD2 was initially characterized as a transcriptional co-activator of estrogen and androgen receptors^9,10^, MYC^3^, and E2F^11^ in human cancer cells, studies of yeast ATAD2 homologs suggest that ATAD2 is more complex, acting not only as a transcriptional regulator but also as a histone chaperone and chromatin boundary element^12^. Moreover, yeast ATAD2 homologs can function either as transcriptional activators or repressors depending on biological context, suggesting that even the transcriptional functions of ATAD2 are poorly understood.

Specifically, the budding yeast ATAD2 homolog Yta7 regulates the expression of many genes including histone gene transcripts by either activating or repressing transcription^13-15^, while the fission yeast ATAD2 homolog, Abo1, generally functions as a transcriptional repressor^16^. Some of the transcriptional roles of Abo1 and Yta7 can be attributed to histone chaperone activities that directly affect nucleosome density and positioning. For example, deletion of Yta7 in budding yeast increases nucleosome density within genes^17^, while in an *in vitro* reconstituted system, catalytically active Yta7 triggers a decrease in histone levels and chromatin disassembly, consistent with a role in gene activation^18^. In contrast, deletion of Abo1 in fission yeast causes a reduction in histone dosage and nucleosome occupancy^16^, while recombinant Abo1 promotes histone assembly in a single-molecule nucleosome assembly assay^19^, suggesting that Abo1 acts as a histone chaperone like Yta7, but instead catalyzes nucleosome assembly instead of disassembly^16^. Besides direct roles in nucleosome assembly and disassembly, Yta7 and Abo1 also act as boundary elements that restrict the spread of silencing at promoters^13,14,20^ and heterochromatin/euchromatin boundaries^21,22^, and show interactions with other histone chaperones such as Rtt106^13,14,20^, FACT^13,16,22^, and Scm3^23^. Furthermore, Yta7 promotes chromatin replication in an *in vitro* reconstituted replication system, suggesting that Yta7 is also involved in replication^18^.

Building on such discoveries from yeast, studies in mammalian cells are beginning to uncover the molecular relationship between human ATAD2 and chromatin. Consistent with roles as a histone chaperone, ATAD2 functions as a generalist facilitator of chromatin dynamics in human ES cells^24^, and also opposes the activity of the HIRA and FACT histone chaperones by limiting their residence times on chromatin^25^. The potential importance of ATAD2 in chromatin replication has also been explored by a study showing the recruitment of ATAD2 to replication sites during S phase^26^.

On a structural level, ATAD2 is a conserved multidomain protein comprised of an N-terminal acidic domain, two AAA+ ATPase domains, a bromodomain, C-terminal domain (Fig. 1a), and an exceptionally long C-terminal linker (CTL) that bridges the bromo- and C-terminal domain. In human ATAD2, only the structure of the bromodomain has been solved experimentally, showing a canonical four-helical bundle structure (consisting of ɑZ, ɑA, ɑB, and ɑC helices) with the connecting ZA and BC loops forming a hydrophobic pocket that coordinates acetyl-lysines of histone tails ^27,28^. This structure explains the binding preference of ATAD2 for acetylated histones and has been vital in the development of currently available ATAD2 inhibitors, but constitutes only a small portion of the full protein.

**Fig. 1.**
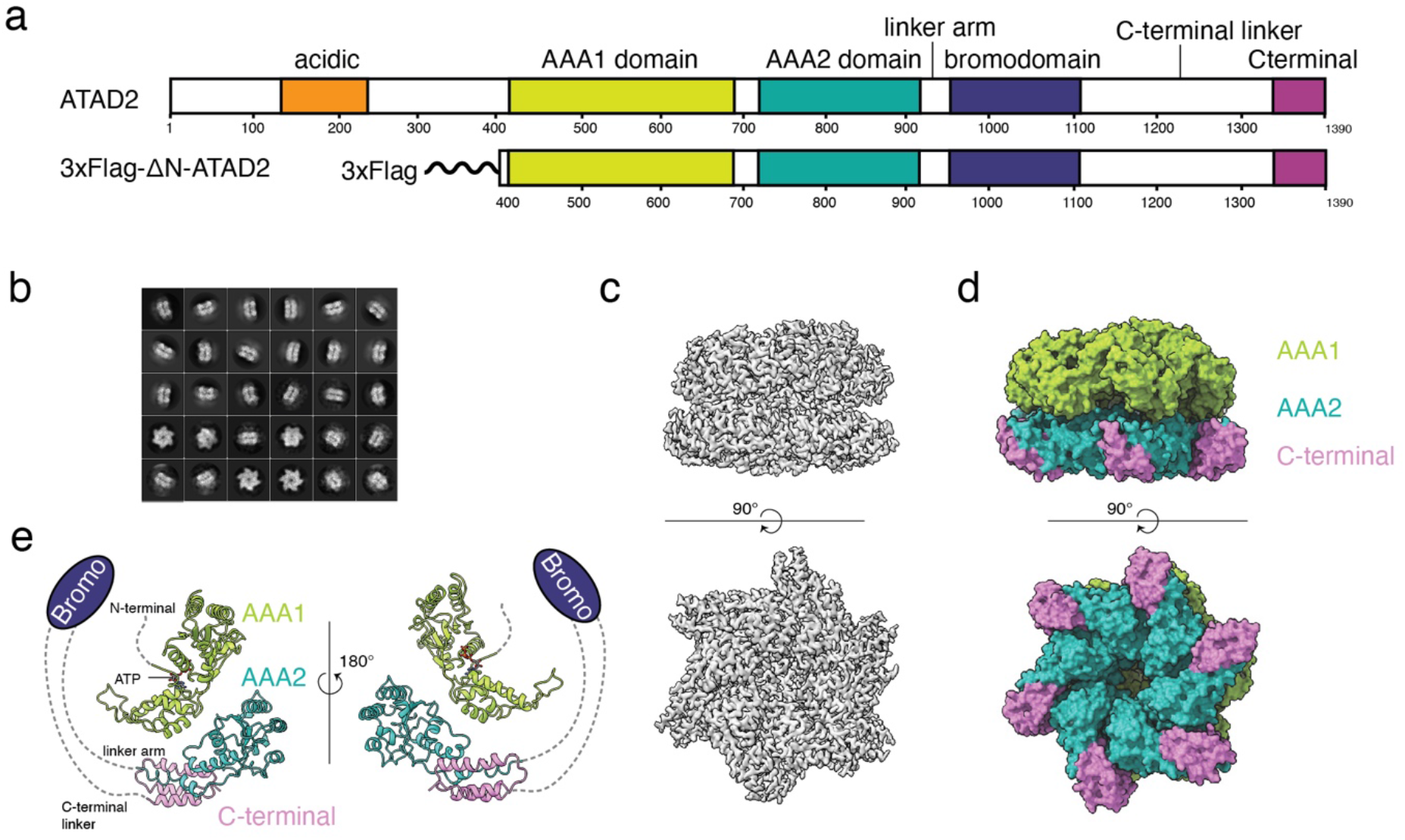
Cryo-EM structure determination of ATAD2. **a** Primary structure of full-length ATAD2 with major domains labeled (top), and the tagged and truncated ATAD2 construct used for structure determination (bottom). **b** 2D class averages of ATAD2 Walker B mutant. **c** Final sharpened density map of ATAD2-Walker B mutant showing side (top) and bottom (bottom) views of the ATAD2 hexamer. **d** ATAD2 Walker B density map color coded by domain as in Fig. 1a. **e** Structure of ATAD2 monomer (chain A) with bound nucleotide, colored by domain.

The distinguishing feature of ATAD2 compared to other histone chaperones is the presence of an ATP-dependent AAA+ ATPase domain. The AAA+ domain is a conserved structural fold shared by the AAA+ ATPase superfamily, where a monomer consists of a large **α**/**β** nucleotide binding domain (NBD), and a small **α** -helical bundle domain (HBD). These monomeric units oligomerize into hexameric rings, where ATP binding pockets are formed at the interface of subunits, and structural motifs from both subunits such as the Walker A/B, sensor I/II, and arginine finger motifs contribute to nucleotide binding and hydrolysis as reviewed in ^29-31^. AAA+ ATPases share common structural and mechanistic features, including dynamic symmetry breaking of rings to spirals, binding and translocation of substrates at the central pore, and the coordinated movement of AAA+ domains with substrate binding domains. Despite similarities in general structural organization, the detailed mechanisms of AAA+ ATPases diverge depending on variations in the structural core, substrate binding domains, and biological functions^32^, and different AAA+ ATPases have evolved different strategies to coordinate the activities of individual subunits within the AAA+ ring.

Our recent structures of Abo1 provide a first look into the overall organization of an ATAD2 homolog, showing that Abo1 assumes a three-tiered hexameric ring structure where a bromodomain ring stacks on top of 2 AAA+ ATPase rings^19^. In these structures, Abo1 AAA+ domains undergo dynamic nucleotide-dependent changes between symmetric rings and asymmetric spirals that alter central pore size and bromodomain arrangement, thus potentially regulating how ATAD2 binds histone substrates. We also find that Abo1 binds the histone H3 tail at the AAA+ ring central pore, which is crucial for Abo1-dependent nucleosome assembly. Despite such advances in understanding Abo1 structural mechanism, whether the divergent human ATAD2 homolog shares common structural features with Abo1 is an open question.

Here, we present cryo-EM structures of near full-length human ATAD2 alone and in complex with the histone substrate H3/H4. We identify an N-terminal linker domain (LD) that wedges between AAA+ subunits at the hexameric spiral seam and potentially locks ATAD2 in a stable conformation. Comparison of the structures of ATAD2 and the ATAD2-H3/H4 complex suggests that the N-terminal LD is undocked by histone H3/H4 binding, potentially unleashing ATAD2 activity and nucleotide-dependent dynamics. Furthermore, we also identify a gate loop that mediates inter-subunit communication in ATAD2, in parallel to inter-subunit signaling (ISS) motifs that have been discovered in other AAA+ ATPases. These results provide the first structural insights into human ATAD2, suggest a unique auto-regulatory mechanism for a AAA+ ATPase, and lay a structural foundation to develop cancer therapeutics.

## RESULTS

### Cryo-EM structure determination of human ATAD2

In our previous studies of Abo1, we found that the acidic N-terminal domain was non-essential for function and overall protein stability^19^. Thus, in this study, we used a truncated version of ATAD2 for cryo-EM structure determination where the acidic N-terminal domain was excluded (Fig. 1a). Unlike Abo1, which forms stable hexamers regardless of nucleotide condition, wild type ATAD2 shows a wide distribution of oligomeric assemblies both in the absence and presence of ATP, as judged by gel filtration and negative stain electron microscopy (Supplementary figs. 1a and b). In addition, wild type ATAD2 does not have measurable ATPase activity, suggesting different characteristics of ATAD2 compared to Abo1. Stable hexamers of ATAD2 are only observed when introducing a Walker B motif (E532Q) mutation that specifically blocks ATP-hydrolysis and mimics AAA+ ATPases in an ATP-bound state (Supplementary figs. 1a and b).

For cryo-EM studies, ATAD2 Walker B mutant protein was purified by sequential affinity, ion exchange, and size exclusion chromatography, and the peak size exclusion fractions corresponding to hexameric size (∼670kDa) were vitrified on cryo-EM grids. Cryo-EM micrographs were collected using a Titan Krios 300 keV with a Gatan Bioquantum GIF/K3 detector, and image processing was performed with Relion 3.1. In short, approximately 2 million particles were picked and subjected to multiple rounds of 2D class averaging (Fig. 1b) and 3D classification. The final cryo-EM map was generated with 212,925 particles to a resolution of 3.15 Å using the 0.143 FSC cutoff criteria (Fig. 1c, Table1, and Supplementary fig. 2). The cryo-EM map shows well-resolved side chains (Supplementary fig. 3a) and reveals an overall structure of two stacked hexameric spirals. Atomic models of six AAA1 domains are built *de novo* into the top-tier spiral, while six AAA2 and C-terminal domains are shown to occupy the bottom-tier spiral (Figs. 1d and e). No density for bromodomains, linker arms, and C-terminal linker domains are visible, likely due to high disorder and/or flexibility of these domains with respect to the AAA domains.

**TABLE 1.**
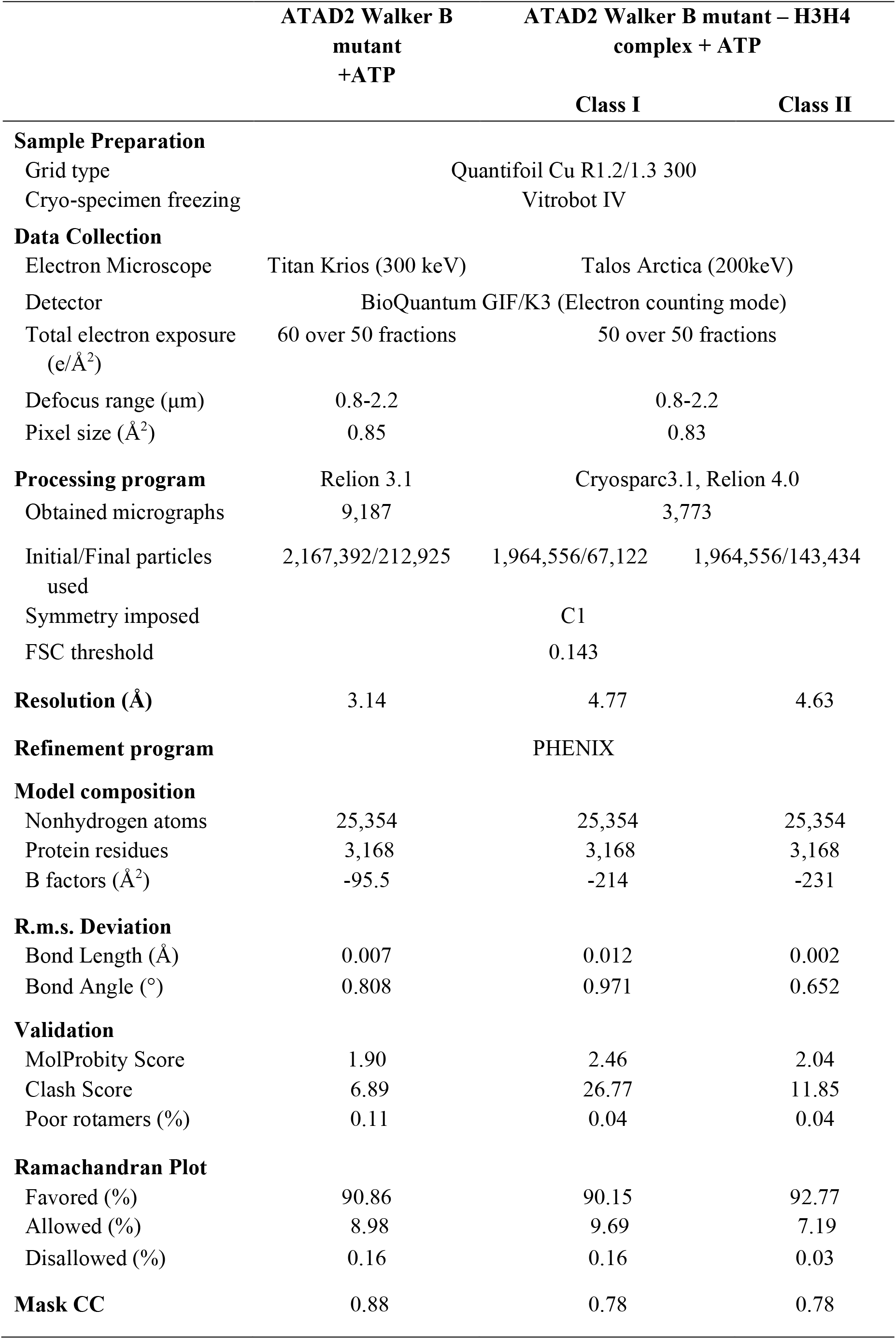
Data collection and refinement statistics.

### Hexameric assembly of ATAD2 compared with Abo1

In the hexameric structure of ATAD2, six subunits are arranged into a right-handed spiral with a rise of ∼2 Å and a rotation of ∼60 ° around the helical axis, resulting in a “seam” between the top (herein referred to as “subunit P1”) and bottom (herein referred to as “subunit P6”) subunits with an offset of ∼12 Å. The hexameric assembly is stabilized by packing of the small ɑ-helical bundle domain (HBD) of one subunit against the large ɑ/β nucleotide binding domain (NBD) of the adjacent subunit, with the nucleotide pocket formed at the interface of adjacent protomers, showing the preservation of the nucleotide binding pocket in ATAD2 as in other AAA+ ATPases^30^. Although the general spiral arrangement of ATAD2 subunits resembles that of Abo1 and other AAA+ ATPase Walker B mutants in an ATP state, ATAD2 has an unusually shallow rise and a narrow seam resulting in a relatively closed and planar spiral (Figs. 2a and b). In addition, in contrast to Abo1 and other AAA+ proteins where the seam subunits are highly disordered^19,33-35^, the P1 and P6 subunits of ATAD2 are well-ordered and have indistinguishable density compared to other non-seam subunits in the local resolution map (Supplementary fig. 3b).

In fact, ATAD2-Walker B-ATP is more similar to the symmetric planar ring conformation of Abo1-ADP (Fig. 2c) or Abo1-apo rather than Abo1-Walker B-ATP, especially in the bottom AAA2/C-terminal ring. The AAA2 domains of P1-P6 do not show a prominent asymmetry or seam (Supplementary fig. 4a), suggesting that the two AAA domain rings are conformationally uncoupled and that movements in the AAA1 domain ring and the AAA1-AAA2 linker are mainly responsible for the helical rise in ATAD2 (Supplementary fig. 4b and c).

**Fig. 2.**
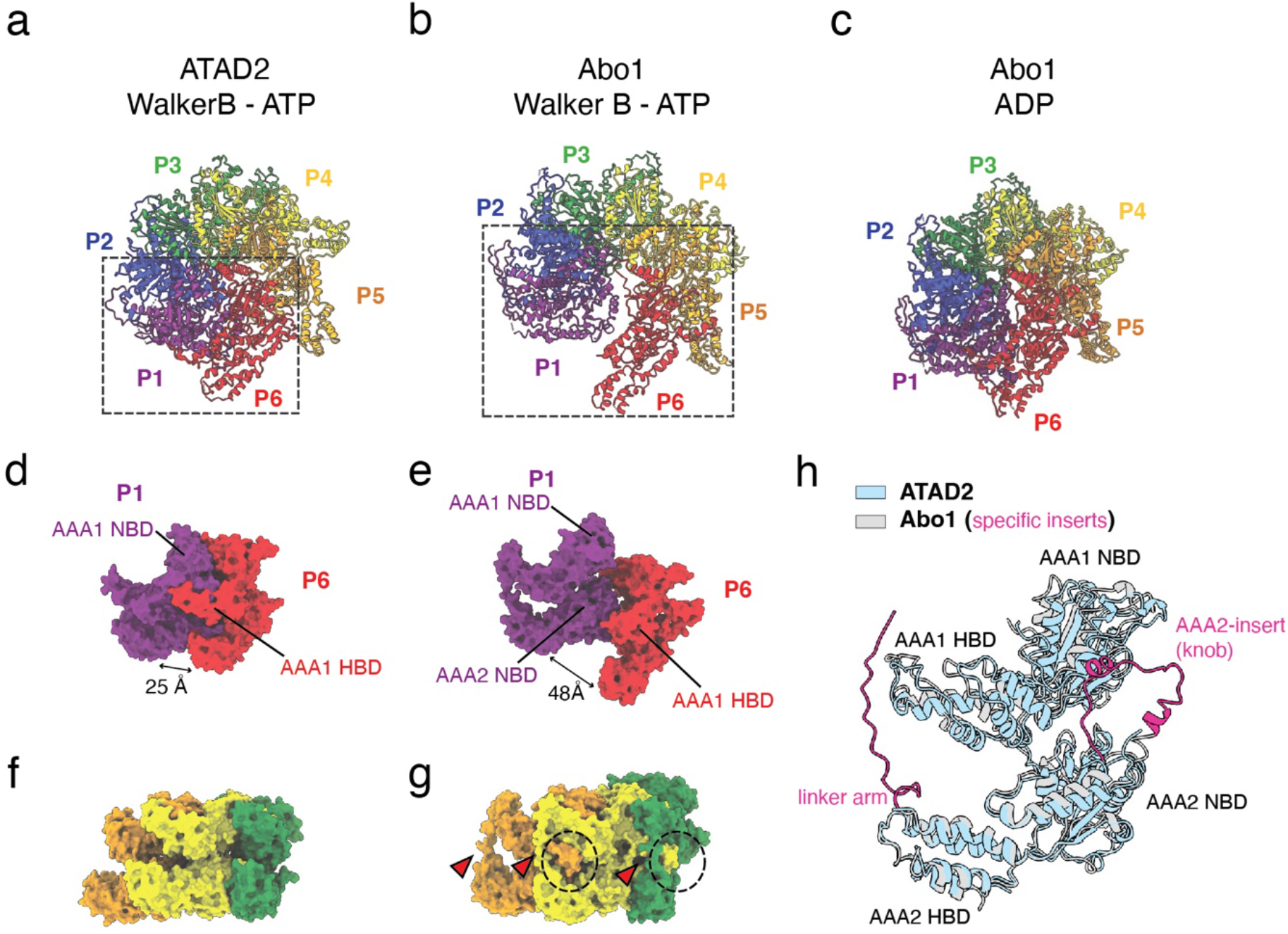
ATAD2 structure in comparison with Abo1. **a-c** Hexameric structures of human ATAD2 Walker B mutant with ATP (**a**), Abo1 Walker B mutant with ATP (**b** PDB ID:6JQ0), and Abo1 with ADP (**c** PDB ID:6JPQ) colored by subunit. Dotted boxes represent spiral seam and are shown as isolated views in panels d-e. **d-e** Seam subunits P1 and P6 from human ATAD2 Walker B mutant (**d**) and Abo1 Walker B mutant (**e**). **f-g** Subunits P3-P5 of ATAD2 Walker B mutant (**f**) and Abo1 Walker B mutant (**g**). Interlocking knob-holes and linker arms are highlighted in (g) with dotted circles, and arrowheads. (**h**) Superimposition of ATAD2 and Abo1 subunits. ATAD2 is shown in light blue and Abo1 in gray. The AAA2-ɑ0 insert knob and linker arm of Abo1 are highlighted in magenta.

When comparing the seam of ATAD2 with Abo1 in the ATP state, it is apparent that the cleft (25 Å vs. 48 Å), and offset (12 Å vs. 35Å) between the P1 and P6 subunits is much smaller in ATAD2 vs Abo1. The difference in offset is such that the P6 subunit AAA1-small helical bundle domain (HBD) packs against the P1 subunit AAA1 large nucleotide binding domain (NBD) in ATAD2, while the P6 AAA1 HBD packs against the P1 AAA2 large NBD in Abo1 (Figs. 2d and e). Further comparison of ATAD2 and Abo1 reveals another key difference. Abo1 has an interlocking knob-hole structure between subunits where a “knob” formed by the AAA2 ɑ0-insert of one subunit inserts into a “hole” formed by the linker arm and small domains of the adjacent subunit^19^, whereas ATAD2 lacks such interlocking structures (Figs. 2f, g, and, h) as the linker arms and ɑ0-insert knobs are disordered in all subunits despite being conserved based on secondary structure alignments. The absence of the knob-hole structure partially explains why in contrast to Abo1, ATAD2 does not form stable hexamers in the absence of nucleotide (Supplementary fig. 1a).

### Nucleotide state and subunit conformation of ATAD2

To better understand the principles of ATAD2 assembly, we examined the structure for the presence of nucleotides and differences in individual subunit conformation. Due to the high resolution of the density map, we were able to ascertain the identity of the nucleotides in all six nucleotide binding pockets of the AAA1 spiral (Figs. 3a and b). Five binding pockets of AAA1 are occupied by ATP, while the binding pocket at the spiral seam (P1/P6 interface) is occupied by ADP. No nucleotides are present in the AAA2/C-terminal spiral, consistent with the absence of key nucleotide binding and hydrolysis sequences in the Walker A and B motifs of AAA2. Although most nucleotide pockets contain ATP, the size of the nucleotide pocket, the orientation of the arginine fingers, and the position of a loop protruding from the adjacent subunit vary among subunits (Fig. 3b).

Further analysis of the individual subunits of ATAD2 (P1-P6) shows that the AAA2/C-terminal domain is relatively rigid with minimal divergence among subunits. However, there are significant differences among subunits in the AAA1 domain, where AAA1 moves as a rigid body with respect to AAA2 due to flexibility in the AAA1-AAA2 linker. In consequence, the AAA1 subunits rise in height and rotate counter-clockwise (when viewing from the top of the AAA ring) with respect to the spiral axis (Fig. 3c, Supplementary fig. 4c, and Supplementary Movie 1) when aligned. Besides rigid body movement of the full AAA domain, the helix ɑ3-sheet β4 loop in AAA1 adopts different conformations (Fig. 3b, d, and e), where the loop protrudes out towards the neighboring subunit in subunits P3-P5, while it is retracted in subunits P1, P2, and P6.

**Fig. 3.**
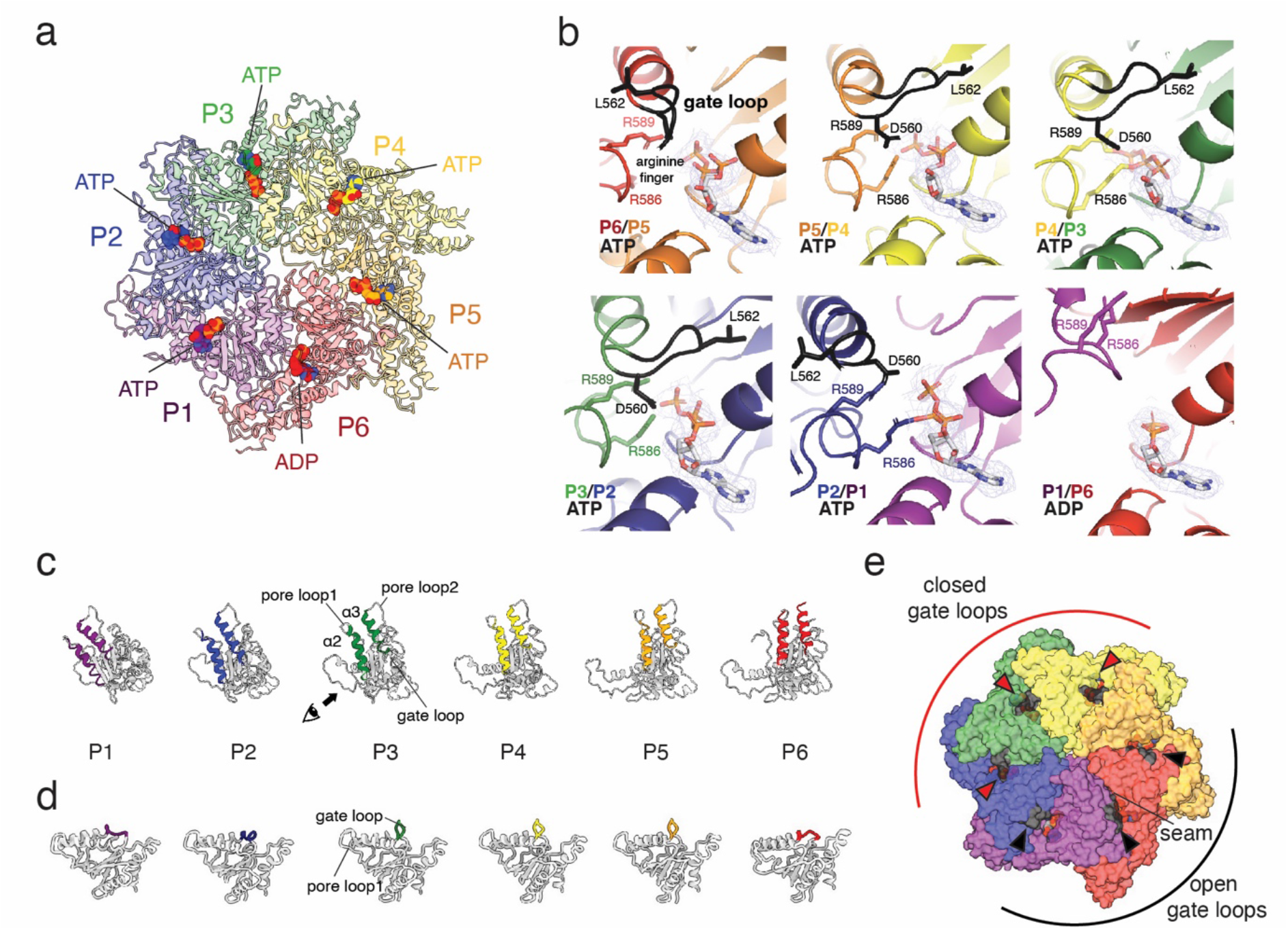
ATAD2 nucleotide pockets and subunits. **a, b** Nucleotide pockets and occupancy of ATAD2. Density map for nucleotide is shown and arginine finger residues R586 and R589 are labeled. ɑ3-β4 gate loops are labeled in black with conserved residues D560 and L562 side chains shown. **c** Variable angles of the AAA1 domain with respect to the AAA2 domain in ATAD2 subunits P1-P6. AAA1 domains are aligned with respect to the AAA2 small domain (AAA2 domains not shown) and shown from a top view of the AAA ring with helix ɑ2 and ɑ3 colored for reference. View from direction of arrow is shown in 1d. **d** Comparison of ɑ3-β4 gate loop conformation in ATAD2 subunits P1-P6. **e** Position of gate loops with respect to nucleotide binding pockets in the ATAD2 hexamer. Gate loops are colored black, with closed gate loops indicated with a red arrowhead, and open gate loops with a black arrowhead.

Interestingly, an equivalent structure termed the “nucleotide communication loop (NCL)” has been observed in the AAA+ ATPase Msp1, where melting of the NCL in response to ATP hydrolysis has been proposed to weaken inter-subunit contacts by disrupting interactions between the NCL and the N-terminal linker domain (LD). The same loop has also been proposed as a nucleotide gate loop in Yta7, closing the nucleotide pocket in the ATP state and opening in the ADP state. Whereas loop conformation directly correlates with nucleotide state in these structures, ATAD2 gate loops assume a distribution independent of nucleotide where the seam-side half of the hexamer has open gate loops and the opposite half of the hexamer has closed gate loops (Fig. 3e). This pattern of open and closed loops is akin to the mixed conformation of inter-subunit signaling (ISS) motifs in a recent structure of p97 where loop conformation reflects the position of a subunit with respect to the post-hydrolysis (ADP or apo) subunit, and the engagement state of the pore loop with a substrate^36^.

Specifically, in the closed gate loops, L562 of inserts into a hydrophobic groove of the adjacent subunit closing off the nucleotide pocket, while D560 stabilizes the position of the two arginine fingers R586 and R589 (Fig. 3b). Although the gate loop of ATAD2 diverges from a conventional ISS motif that was originally defined by a conserved DGF tripeptide constituting part of the ɑ3 helix in membrane-bound AAA+ proteases^37^ (Supplementary fig. 5), the closed conformation shows essentially the same structure as an ISS motif, and likely functions in an identical manner.

### An N-terminal linker domain (LD) that stabilizes the ATAD2 hexamer

When comparing ATAD2 subunits we also discovered a conserved N-terminal linker domain (aa406-423, referred to as “LD” hereafter, Supplementary fig. 6) that was only visible in the ADP-bound subunit, P6, and was sandwiched between the two subunits at the spiral seam (subunits P1 and P6, Fig. 4a). Structural alignments with other AAA+ ATPases revealed that the ATAD2 N-terminal LD assumes the same position as the N-terminal LD of the microtubule severing enzymes katanin^38^ and spastin^39,40^ (termed the “fishhook” element), and the membrane protein extracting enzyme Msp1^35^ (termed the “LD linker domain”) (Fig. 4b). In microtubule severing enzymes and Msp1, this domain consists of two helices (ɑ0 & ɑ1) connected by two linkers (L1 &L2) curved into a “fishhook-like” shape, and lies close to the initial substrate binding surface on top of the spiral. Thus, this structural domain has been proposed to mediate inter-subunit communication of nucleotide and substrate binding state.

**Fig. 4.**
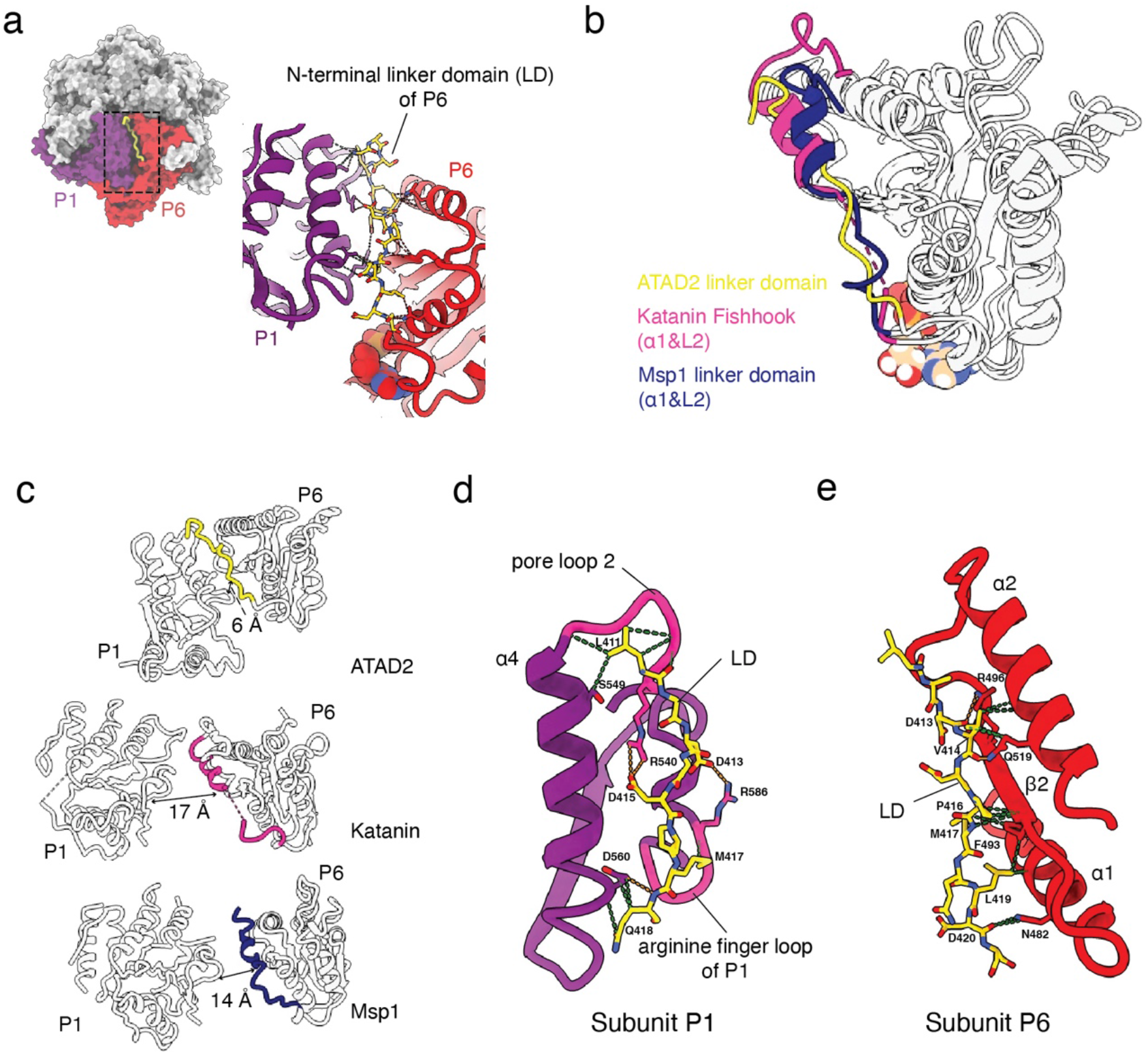
Structure of the ATAD2 N-terminal linker domain (LD) **a** Position of the ATAD2 subunit P1 N-terminal LD (yellow) wedged between subunit P1 (purple) and P6 (red) in the hexameric spiral. **b** Superimposition of the large nucleotide binding domains of ATAD2, katanin (PDB ID: 6UGD) and Msp1 (PDB ID: 6PE0) highlighting the position of the LD with respect to the nucleotide binding domain body in yellow, hotpink, and blue, respectively. **c** Comparison of distance from P6 LD to adjacent P1 subunit in ATAD2, Katanin, and Msp1. Labeled distances are measured from middle of P6 LD ɑ1 helix to P1 arginine finger. **d**,**e** Detailed hydrogen (orange) and van der waals interactions (green) of the ATAD2 P6 LD with (**d**) the P1 subunit ɑ4 helix, pore loop 2, and arginine finger loop (pore loop 2 and arginine finger loop highlighted in magenta), and (**e**) the P6 subunit ɑ1, ɑ2 helix, and β2 sheet.

In the ATAD2 structure, the N-terminal LD consists of a loop that corresponds to ɑ1 and L2 of the katanin fishhook or Msp1 LD. Further comparison of N-terminal LD interactions with adjacent domains reveals parallels between katanin and ATAD2 (Fig. 4c). In both structures, the N-terminal linker interacts with the pore loop of the clockwise subunit, as well as helix ɑ2 of the same subunit. However, the ATAD2 N-terminal LD has much more extensive interactions that are absent in Msp1 and microtubule severing enzymes, such as interactions with the arginine finger loop in the adjacent P1 subunit, and with helix ɑ2 and sheet β1 in the same P6 subunit (Figs. 4d and e). The tight cleft is also illustrated by comparing the distance from the N-terminal linker to the arginine finger loop of the adjacent subunit, where the distance is only 6 Å in ATAD2 compared to 17 Å and 14 Å in katanin and Msp1, respectively.

Our structure shows that D415 from the N-terminal LD of P6 and R540 from P1 pore loop 2 form a salt bridge which may stabilize the P1 N-terminal LD-P6 interaction and the hexamerization of the ATAD2. To test this, we engineered D415A/R540A mutations in the Walker B mutant (E532Q) background. In contrast to the Walker B mutant, the LD/Walker B triple mutant (D415A/E532Q/R540A) displayed a broader size distribution with heterogeneous particles despite high expression levels and purity (Supplementary fig. 1).

Thus, the N-terminal LD seems to effectively lock the ATAD2 hexamer in a stable conformation, where ATAD2 is unable to trigger ATP hydrolysis and symmetry breaking at another subunit. Consistent with this idea, the seam subunits of ATAD2 are well-ordered with resolution similar to other non-seam subunits (Supplementary fig. 3b), in contrast to other AAA+ proteins where the seam subunits are disordered, and the bottom subunit of the spiral usually diverges significantly^33^. Although we were unable to determine the functional effects of LD mutations on ATAD2 as a structurally homogeneous and catalytically active form of wild type ATAD2 was unattainable, we predict that disruption of the D415-R540 salt bridge might increase basal ATPase rates in wild type ATAD2.

### Pore loop and substrate binding of ATAD2

After modelling all six subunits of ATAD2 into the density map, we also noticed a small region of extra density in the central pore of AAA1. This extra density was likely part of a substrate that co-purified with ATAD2 from Sf9 insect cells, as was observed in Abo1^19^. Side chains of the substrate were undiscernible, so we modelled the substrate as a linear peptide of 5 alanines with the N-terminus facing toward the interior of the pore, consistent with other AAA+ ATPases (Fig. 5a, left). The AAA1 pore loop1 of ATAD2 wrapped around the substrate with a spiral staircase (Fig.5a, right) that has been shown to be structurally well conserved in various AAA+ ATPases ^33^. Consistent with this, ATAD2 AAA1 pore loop 1 W505 residues engage the peptide substrate by inserting an orthogonally positioned aromatic ring between two side chains of the substrate. However, compared to other AAA+ structures and even the homologous Abo1 structure, ATAD2 shows signs of a weakly bound peptide.

**Fig. 5.**
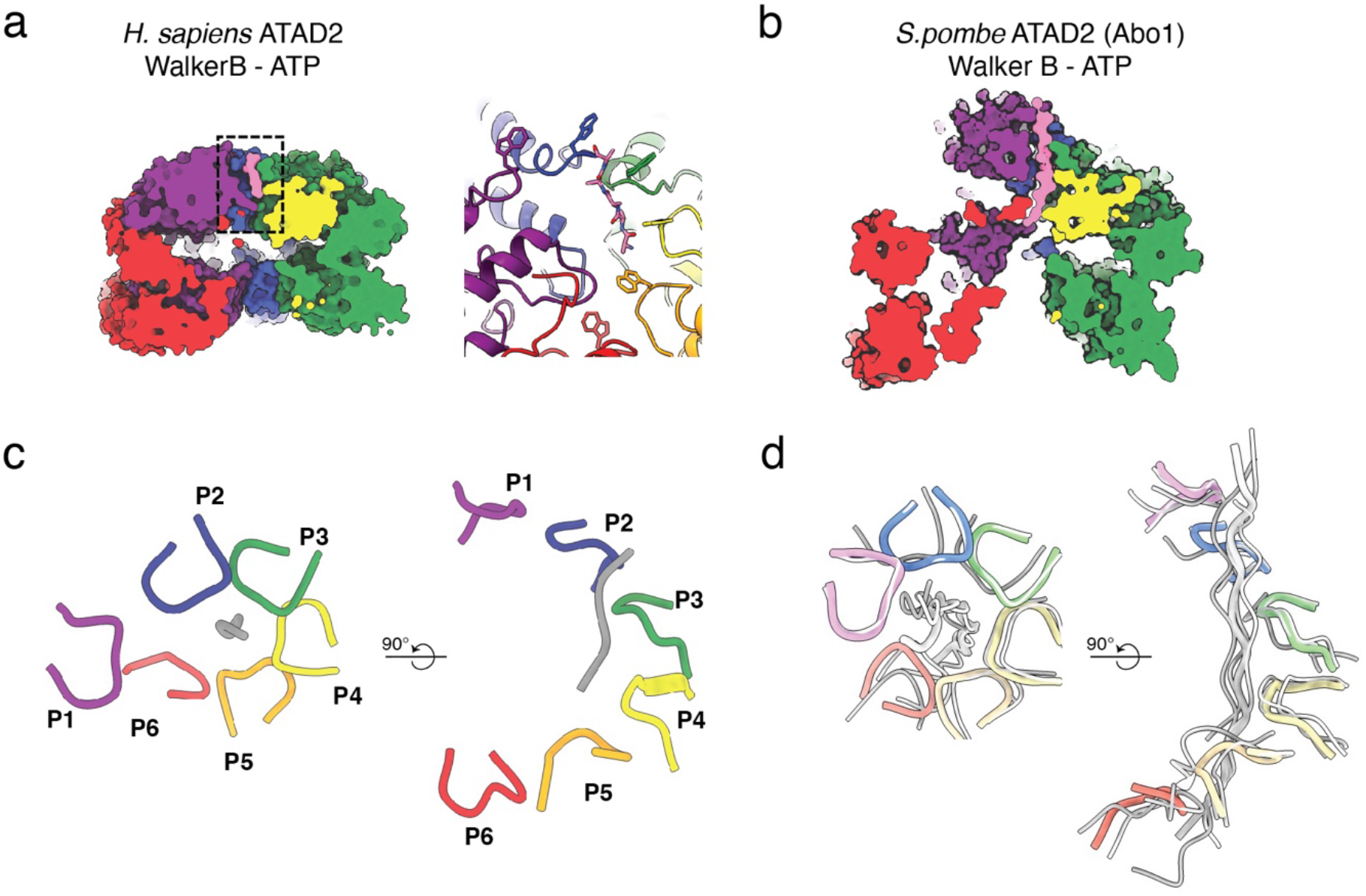
ATAD2 substrate binding at the AAA+ pore. **a**,**b** Cut-open side view showing a substrate (pink) bound to the pore of ATAD2 (**a**), and Abo1 (**b**). Boxed region in (**a**) is shown in detail at right showing ATAD2 pore loop staircase with W505A surrounding a peptide substrate. **c** Top (left) and side (right) views of ATAD2 pore loop (colored by chain) - substrate (gray) interactions. **d** Top (left) and side (right) views of superimposed pore loop-substrate interactions of Abo1 (colored by chain, PDB ID: 6JQ0), spastin (grey, PDB ID: 6PEN), and NSF (grey, PDB ID:6MDO).

First, the peptide substrate shows a weak density where only 5 aa can be modelled, as opposed to 14 aa or 22aa substrates in Abo1or translocating unfoldases p97. Thus, in contrast to Abo1 where a long stretch of peptide is inserted deep into the asymmetric spiral, only a short peptide is docked shallowly near the top surface of ATAD2 (Figs. 5a and b). Second, only AAA1 pore loop1 grips the peptide, while AAA1 pore loop 2 and the pore loops of AAA2 do not participate in substrate interaction. Third, while translocating AAA+ ATPases usually have 5 to 6 subunits engaged with the substrate in the active state, only 3 ATAD2 subunits (P2-4) engage with substrate, suggesting a “weak grip”, with the P1 and P6 subunits displaced significantly from the conventional pore loop staircase (Figs. 5c and d). Notably, the subunits that participate in substrate interactions correspond to the subunits with nucleotide pockets closed off by gate loops (Fig. 3e).

### Structure of the ATAD2-histone H3/H4 complex

To understand how ATAD2 acts on its substrate, we mapped the interactions of ATAD2 Walker B mutant with recombinant histone H3/H4 by crosslinking mass spectrometry (XL-MS) (Supplementary fig. 7). ATAD2 crosslinks with histone H3/H4 at multiple positions through the AAA1, bromo- and C-terminal linker domains, while the AAA2 and C-terminal domains do not show any crosslinks. This is in agreement with the overall architecture of ATAD2, where the AAA1 and bromodomains form the top histone interacting surface, while the AAA and C-terminal domains form the inert base on the bottom. Moreover, all AAA1 residues crosslinking to histone H3/H4 map to the top surface of the AAA ring, further supporting the idea that the histone binding surface is formed by the top surface of the AAA1 ring, bromodomains, and C-terminal linker domains. Multiple positions on histone H3/H4 also crosslink to ATAD2, both within the N-terminal tails and histone bodies. Akin to Abo1, the N-terminus of H3 crosslinks to AAA1 pore loop 1 suggesting that the substrate observed in the ATAD2 structure is also part of the histone H3 N-terminal tail.

Further cryo-EM structure determination of the crosslinked ATAD2 Walker B mutant-histone H3/H4 complex reveals two conformations that have extra density on top of the AAA1 ring (Figs. 6a and b, Supplementary fig. 8, and Table 1). In the first conformation (Class I, 4.7Å resolution), the extra density shows a helical handle-like structure that connects to the P1 subunit (Fig. 6a). By rigid body docking, two copies of the human ATAD2 bromodomain (PDB ID: 3DAI) fit into the extra density in a head-to-tail arrangement, which is consistent with the orientation of the bromodomains observed in Yta7 (Fig. 6c) ^41^.

**FIGURE 6.**
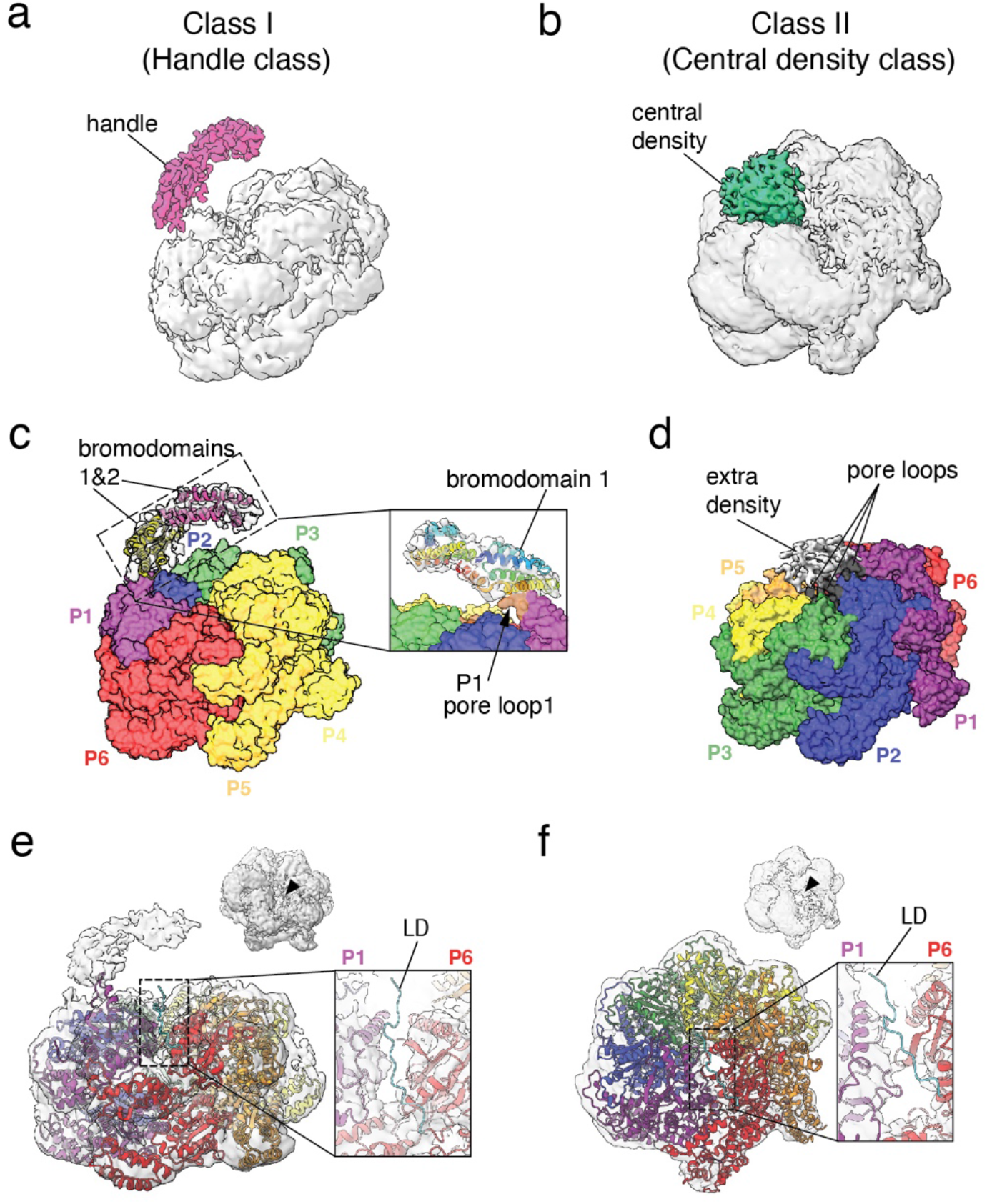
**a**,**b** Density maps of ATAD2 Walker B mutant-H3H4 complex from class I (**a**) and II (**b**) with extra density after subtracting AAA+ domains colored hot pink (**a**) and sea green (**b**), respectively. **c** Class I model with surface representation of 6 AAA+ domains colored by subunit and two bromodomains (bromodomain 1 in olive, bromodomain 2 in hot pink) docked into extra density. Inset shows enlarged view of boxed region showing potential contact between AAA+ ring and bromodomain where P1 subunit pore loop 1 (brown) contacts bromodomain 1. **d** Class II model with surface representation of 6 AAA+ domains colored by subunit and extra density at center. Pore loop 1 of all subunits are colored black. **e**,**f** Class I (**e**) and class II (**f**) density maps with fitted AAA+ domains showing a view of cleft at the P1 (purple) /P6 (red) subunit seam. Inset shows enlarged view of cleft with LD highlighted in cyan. Black arrowheads indicate position of seam.

Interestingly, in this state, the bottom of bromodomain 1 seems to make potential contacts with the P1 subunit pore loop 1. In the second conformation (Class II, 4.6 Å resolution, Fig. 6b), an extra globular density is observed in the central pore, and has a broad interface with the AAA1 domains of P1-P3 with the pore loops lining the bottom surface (Fig. 6d). The resolution of the extra density is insufficient to unambiguously determine the identity of the substrate, but based on the volume of the extra density and previous structures of ATAD2 homologs^19,41^ the most probable candidate would be a histone H3/H4 dimer.

In order to understand conformational changes in the AAA+ ring induced by histone H3/H4 binding, the structure of ATAD2 alone was fit and refined into the ATAD2-histone H3/H4 complex maps. In agreement with XL-MS data that shows no major differences between the crosslinking patterns of ATAD2 and ATAD2-histone H3/H4, the ATAD2 structure fit well into both class I and class II maps without any major rearrangements, suggesting that ATAD2 does not go through large-scale conformational changes upon H3H4 binding.

However, there is a notable difference between ATAD2 and ATAD2-histone H3/H4 maps at the seam (P1/P6), where the P6 subunit is slightly disordered in class I, and more prominently disordered in class II (Fig. 6e, f, and Supplementary fig. 9). As a result of the cleft of missing density, the N-terminal LD of the P6 subunit cannot be placed in the density map indicating that it may be undocked from its original position sandwiched between P1 and P6. Consistent with this, additional examination of ATAD2 intramolecular crosslinks shows that the N-terminal LD crosslinks extensively with the nucleotide binding pocket, pore loop 1, bromodomain, and C-terminal linker (CTL) domain in the absence of histone H3H4, but loses most of these crosslinks upon binding histone H3/H4 (Supplementary fig. 7).

Furthermore, superimposition of the AAA+ domains from ATAD2 and ATAD2-histone complexes shows that in the ATAD2-histone complexes, movement of the P1 and P6 subunits away from each other widens the seam cleft, and overall opens the AAA ring (Supplementary Movies 2 and 3). Together, these data support the idea that H3H4 binding to the bromo- and CTL domain might allosterically release the N-terminal LD from the P1/P6 seam, opening up the AAA+ ring, and priming the AAA+ ring for activation.

## DISCUSSION

### An autoinhibited state of ATAD2

In this study we present the atomic resolution cryo-EM structure of the human ATAD2 AAA+ ATPase, showing an architecture that follows the general assembly principles of AAA+ protein unfolding and disassembly machines. ATAD2 forms a shallow hexameric spiral staircase that binds a peptide, potentially the histone H3 tail, by conserved aromatic loops at the central pore.

Several lines of evidence suggest that we have captured ATAD2 in a special autoinhibited state that has been previously unobserved in other AAA+ ATPases. First, in many AAA+ ATPases, breaking of ring symmetry to spirals has been shown to be essential for mechanical translocation of substrates and disassembly activity. Thus, subunits at spiral seams are disordered and variable in position as they have been caught in the act of translocation. In contrast, our ATAD2 structure has well-ordered subunits at the spiral seam that have indistinguishable resolution from other subunits (Supplementary fig. 3a) suggesting limited mobility and symmetry breaking. Second, we observe the N-terminal LD of the P6 subunit wedged between the seam subunits, where it effectively glues the P1 and P6 subunits in place, thus limiting subunit translocation and symmetry breaking. Third, only three ATAD2 pore loops maintain a weak grip on a short length of substrate in the central substrate, hinting at a non-productive or weakly-productive state if ATAD2 were to pull on substrates. Together, these lines of evidence imply that the N-terminal LD acts as a brake that maintains ATAD2 in an autoinhibited state thus preventing premature ATPase activation.

Interestingly, the N-terminal LD was also proposed as a gate loop in recent structures of the ATAD2 yeast homolog, Yta7 based on its nucleotide-dependent conformational change near the nucleotide entry pocket^41^. Although the gate loop in Yta7 is not docked to seam subunits as in ATAD2 (discussed in more detail below), the idea of the N-terminal LD as a gating element that regulates nucleotide hydrolysis and exchange might be conserved between species.

### Binding of ATAD2-H3H4

Our structure of the ATAD2-histone H3H4 complex shows the N-terminal LD displaced from its original position and the ATAD2 hexamer with more disorder at the seam, suggesting that the binding of histone H3/H4 to ATAD2 somehow undocks the LD through allosteric changes. Further clues to how histone H3/H4 binding might induce conformational changes in ATAD2 come from XL-MS. The N-terminal LD crosslinks not only with the AAA NBD as predicted by the cryo-EM structure, but also with the bromo- and C-terminal linker domains that are invisible in the cryo-EM structure. Many of these intramolecular crosslinks disappear in the histone H3/H4-bound complex, and are replaced with intermolecular crosslinks to histone H3/H4. Thus, it is plausible that the N-terminal LD serves as a major integration point where information on histone substrate binding is communicated to the AAA+ domain, and is also consistent with the role of the LD proposed in other AAA+ ATPases such as Msp1. Such a mechanism might be required to keep ATAD2 enzyme activity in check until recognition of an H3H4 substrate.

Besides suggesting a mode of ATAD2 activation, the ATAD2-histone H3/H4 complex structure also provides insight into how ATAD2 binds its substrate. In the class I conformation of the ATAD2-histone complex, two bromodomains form the base of a spiral that protrudes up from the top surface of the ATAD2 ring. Although density for histone substrate is not directly observed in this conformation, this is possibly a primed state that is conducive for initial weak binding of histones by the bromodomains as the N-terminal LD is undocked and AAA ring symmetry is broken. It is also interesting to note that AAA1 pore loop forms putative contacts with the bromodomains in this conformation, which would be incompatible with H3 peptide binding at the pore and is indicative of a weak substrate binding state where histones are not completely docked. In the class II conformation, the substrate, potentially histone H3/H4, is embedded by a larger surface area at the central pore with at least 3 subunits and pore loops engaged. This conformation also has more disorder and missing density at seam subunits suggesting that stable binding and docking of H3H4 to the AAA ring central pore further breaks symmetry and activates ATAD2.

Although we observed unmodified histone binding to ATAD2 in this study, it will be interesting to investigate how acetylated histone H3/H4 alters ATAD2 conformation in future studies as the ATAD2 bromodomain has been shown to prefer H4 acetylated at K5, K8, K12, and K16^42,43^.

### Comparison of ATAD2 homologs and implications for function

Two major structural differences between human ATAD2 and its yeast homologs emerge from this study, which might allude to differences in roles and mechanisms. First, the absence of interlocking knob-hole interactions in ATAD2 compared to Abo1 and Yta7 suggests that human ATAD2 forms a more dynamic and labile hexamer that can disassemble into monomers more readily. This is consistent with our biochemical results showing that wild type ATAD2 assumes a wide range of oligomeric forms in contrast to Abo1, which is a stable hexamer irrespective of nucleotide condition. Second, the N-terminal LD wedges between subunits in the ATAD2 structure, but is not observed in the Abo1 or Yta7 structures. Although the LD may play a nucleotide gating role in all homologs, divergence in N-terminal LD and the interacting bromo- and CTL domains, as well as differences in hexameric organization may explain why the LD acts as a brake only in ATAD2. Together, these differences might reflect the different substrate binding preferences (ATAD2 prefers acetylated histones while Abo1 and Yta7 have no preference) and functions (promotion of nucleosome disassembly vs. assembly) of different homologs.

### An inter-subunit signaling (ISS) motif in ATAD2

Another important finding of this study is the discovery of a gate loop that in ATAD2 that corresponds to ISS motifs in other AAA+ ATPases. ISS motifs were discovered in classical clade m-AAA proteases as elements that transmit information about the nucleotide state to adjacent subunits and pore loops^37^, and were originally defined by the presence of a crucial phenylalanine in a conserved DGF tripeptide constituting part of the ɑ3 helix in the AAA NBD ^37^. However, ISS motifs can diverge with a short insertion positioned C-terminal to the traditional ISS motif, such that in some cases like Msp1, it has even been termed a different name (the “nucleotide communication loop (NCL)”). Sequence -wise, the ATAD2 ɑ3-β4 loop diverges from the originally discovered ISS motif, but shows similar conformations to the p97 ISS, where the loops jut out in a triangular shape towards the neighboring nucleotide binding pocket. Thus, it seems that the category of ISS motifs can be expanded to include a wider variety of sequences in the ɑ3 C-terminus and ɑ3-β4 loop than originally defined (Supplementary fig. 5).

### ATAD2 function and pharmacological implications

Despite advances in understanding ATAD2 structure in this study, we were unable to purify a catalytically active from of wild type ATAD2 despite various attempts with different constructs and conditions. Thus, for now, a direct demonstration of ATAD2 function *in vitro* awaits testing. However, we have paved the road to design inhibitors targeting the ATAD2 AAA+ domain by revealing two gating elements: the N-terminal LD and an ISS-like motif. Considering that recent structures of the AAA+ ATPase p97 show the p97 inhibitor NMS-973 binding to the ISS motif,^36^ and that ISS loops have variable sequences in different AAA+ ATPases, our structure offers promising new strategies to target ATAD2.

## METHODS

### Cloning and protein overexpression

DNA encoding aa 403-1390 of ATAD2 was PCR amplified from the full-length codon-optimized synthetic ATAD2 gene (Genscript), and subcloned into an in-house modified pFastBac1 vector with an N-terminal 3xFlag tag and Tev protease cleavage site using EcoRI and XhoI restriction sites. Point mutants of ATAD2 (E532Q, D415A, and R540A) were created by inverse PCR with KOD plus polymerase (Toyobo) according to product manual instructions. Cloned plasmids were transformed into DH10Bac competent cells to produce bacmids that were subsequently transfected into SF9 cells to produce baculovirus, and baculovirus was amplified to P3 according the Bac-to-Bac Baclulovirus Expression Kit (ThermoFisher Scientific). ATAD2 proteins were expressed by infecting 950mL of SF9 suspension cultures at a density of 2.5-3.5 × 10^6^ cells with 50mL of P3 virus for 45-48hrs. Recombinant *X. laevis* histones H3 and H4 were overexpressed in *E. coli* and purified by inclusion body purification as described in Luger et al ^44^.

### Protein purification of ATAD2 and H3H4

Harvested SF9 cells were resuspended in 20mL of lysis buffer (50mM Tris-HCl (8.0), 300mM NaCl, 5% glycerol) per liter cells, and lysed by 4 freeze/thaw cycles in the presence of 1mM PMSF and cOmplete protease inhibitor cocktail tablet (Roche). Lysates were clarified by centrifugation at 39,000xg for 1.5hrs, and were batch bound to 1mL of anti-DYKDDDDK G1 affinity resin (Genscript) per L cells for 2hrs. Resin was washed with a sequence of 5 column volumes (CV) lysis buffer, 5 CV wash buffer 1 (50mM Tris-HCl (8.0), 500mM NaCl, 5% glycerol), 5 CV wash buffer 2 (50mM Tris-HCl (8.0), 1000mM NaCl, 5% glycerol), followed by 5 CV wash buffer 1, 5 CV was buffer 2, and 5 CV lysis buffer.

Proteins were eluted by incubating Flag resin with Flag elution buffer 1 CV at a time for 10min (lysis buffer with 0.08mg/mL DYKDDDDK peptide (Shanghai Apeptide)). A total of 5-7CV were eluted and checked for yield and purity by SDS-PAGE. Peak fractions were pooled and diluted with an equal volume of no salt buffer (50mM Tris (8.0), 5% glycerol), and loaded onto a HiTrapQ 5mL column (Cytiva Life Sciences). Fractions were eluted with a 50mM to 1000mM NaCl gradient over 20CV. Peak fractions with a A260/280 ratio of < 0.6 were pooled, concentrated with an Amicon Ultra-15 100kDa cutoff centrifugal concentrator (Millipore), and run over a Superose6 increase 10/300GL (Cytiva Life Sciences) column in 25mM HEPES, 250mM NaCl, 5% glycerol, and 1mM DTT. Gel filtration peak fractions corresponding to hexameric ATAD2 were pooled, concentrated and flash frozen until further use. Typical protein yields were on the order of ∼100ug purified protein per L cells.

Histone H3/H4 complex was prepared as described in Luger et al^44^. Briefly, *X. laevis* histones H3 and H4 were extracted in their denatured forms from bacterial inclusion bodies, and purified further by gel filtration and cation exchange chromatography. Purified H3 and H4 were dissolved in unfolding buffer (20mM Tris-HCl (7.5), 6M Guanidine-HCl, and 5mM DTT) and mixed at equimolar ratios, refolded by dialysis overnight in high salt buffer (10mM Tris-HCL (7.5), 2M NaCl, 1mM EDTA, and 5mM β-mercaptoethoanol), and purified over a Superdex 200 26/60 (Cytiva Life Sciences) column. Peak fractions corresponding to histone H3/H4 tetramer were pooled, concentrated, and used for ATAD2-histone H3/H4 complex formation.

### Preparation of ATAD2-histone H3/H4 complex

To obtain ATAD2-histone H3/H4 complexes for cryo-EM data collection, ATAD2 Walker B mutant protein and refolded histone H3/H4 were mixed at a 1:1.5 molar ratio in a buffer consisting of 25mM HEPES (7.5), 150mM NaCl, 5% glycerol, and 1mM DTT, and incubated for 30 min on ice. To initiate crosslinking, 1mM of DSS (Sigma-Aldrich) was added to proteins and incubated for 30min @ 37 degrees, and quenched by the addition of ammonium bicarbonate to a concentration of 50mM and an additional 20min incubation @ 37 degrees. Crosslinked ATAD2-histone H3/H4 complex was further purified by a 10-30% sucrose gradient run for 17hrs at 65,000xg to remove over-crosslinked aggregates and excess histone H3/H4.

### Cryo-EM grid preparation and data collection

Before vitrification, proteins were buffer exchanged to 25mM HEPES (7.5), 250mM NaCl, 0.025% beta-octyl-glucoside, 1mM DTT, and 1mM Mg-ATP and concentrated to 1.8 mg/mL. Proteins were frozen on glow-discharged Quantifoil Cu R1.2/1.3 300 mesh grids with a Vitrobot IV (Thermo Fisher Scientific) set at 15℃ and 100% humidity and blotted with -10 or 0 blotting force with 3 sec blotting time. For the ATAD2 Walker B -ATP dataset, 9,187 micrographs were collected using a Titan Krios 300 keV microscope with a Gatan Bioquantum GIF/K3 direct detector varying the defocus of 0.8 to 2.2µm with 0.5 µm step at Harvard Center for Cryo-Electron Microscopy (HC2EM) (Table 1 and Supplementary fig. 2). For the ATAD2 Walker B-histone H3/H4-ATP dataset, 3,686 micrographs were collected using a Talos Artica 200kEV microscope with a Gatan Bioquantum GIF/K3 direct detector varying the defocus of 0.8 to 2.2µm with 0.5 µm step at KBSI Ochang.

### Cryo-EM data processing

For the ATAD2 alone dataset, all processing was performed in Relion 3.1^45^. A total of 2,167,392 particles were picked from motion corrected (using MotionCorr2) and CTF corrected micrographs (using CTFFIND4.1) and subjected to multiple rounds of 2D class averaging (Supplementary fig. 2). Multiple rounds of 3D classification starting with the good class averages with 417,689 particles followed by refinement and postprocessing were performed. The final 3.1Å resolution map was obtained based on 0.143 FSC cutoff criteria. An atomic model of ATAD2 was built de novo into the cryo-EM map using COOT followed by PHENIX refinement.

For the ATAD2-histone H3H4 complex dataset, a total of 1,964,556 particles were picked from motion corrected (using MotionCorr2) and CTF corrected micrographs (using CTFFIND4.1). Multiple rounds of 2D class averaging (Supplementary fig. 8) and classification based on ab initio models was performed in cryoSPARC 3.1^46^. 346, 483 particles were moved from cryoSPARC to Relion 4.0 by pyEM, and 3D classification with alignment was performed to yield class I and class II particles. Refinement and postprocessing were performed on class I and class II particles using non-uniform refinement in cryoSPARC yielding final 4.7Å (class I) and 4.6 Å (class II) resolution maps based on 0.143 FSC cutoff criteria. Structures of ATAD2 obtained from ATAD2 cryo-EM maps were fit and refined into ATAD2-H3H4 complex maps, and bromodomains (PDB ID 3DAI) were docked into maps in Phenix. Final cryo-EM maps and structure coordinates are deposited in EMDB (EMD-34468, 34471, and 34472) and PDB (ID: 8H3H, 8H3O, and, 8H3P).

### Crosslinking mass spectrometry (XL-MS)

ATAD2-histone H3/H4 complexes were prepared by the same method as for cryo-EM samples, with the exception that 1mM of DSS H12/D12 (Creative Molecules) was used instead of 1mM DSS. Crosslinked ATAD2-histone H3/H4 complexes were purified on a 10-30% sucrose gradient as with cryo-EM samples and buffer-exchanged to 25mM HEPES (7.5), 250mM NaCl, 5% glycerol, 1mM DTT, and 1mM Mg-ATP. Denaturation, blocking, and digestion of proteins were performed essentially as in^19^. Digested peptides were separated by passage over a Superdex peptide 3.2/30 column, and analyzed on an Orbitrap mass spectrometer (Thermo Fisher Scientific) at the Taplin Mass Spectrometry facility at Harvard Medical School. Data analysis was performed using xQuest^47^ and a sequence database containing ATAD2, histone H3/H4 sequences. xQuest search results were filtered according to the following criteria: mass error < 4 ppm, minimum peptide length = 6 residues, delta score < 0.9% TIC ≥ 0.1, minimum number of bond cleavages per peptides = 4, and an xQuest LD score cutoff of 25 was selected, corresponding to a false discovery rate of < 2%. Final crosslinks were filtered and visualized with xiView^48^.

## Supporting information

Summplementary Information

## ACKNOWLEDGEMENTS

We would like to thank Sarah Sterling and Richard Walsh at the Harvard Center for Cryo-Electron Microscopy (HC2EM), and Dr. Sunghoon Chun at Korea Basic Science Institute (KBSI, Ochang) for supporting data collection. We also thank Yumi Shin and Eunhee Seong for technical assistance, and Dr. Bob Kingston for supporting this project. This work was supported by grants from the National Research Foundation of Korea (NRF). Specifically, a Sejong Science Fellowship (2022R1C1C2003419) and the Basic Science Research Program through the Korea Ministry of Education (2019R1A6A1A10073887) to C.C. and grants (2020R1A2B5B03001517, 2020M3E5E2037170) and the framework of international cooperation program (2021K2A9A2A08000088) to J.S..

## CONFLICTS OF INTEREST

J.S. is a co-founder and CTO of Epinogen.

## REFERENCES

1. Caron, C. et al. Functional characterization of ATAD2 as a new cancer/testis factor and a predictor of poor prognosis in breast and lung cancers. Oncogene 29, 5171–81 (2010).

2. Boussouar, F., Jamshidikia, M., Morozumi, Y., Rousseaux, S. & Khochbin, S. Malignant genome reprogramming by ATAD2. Biochim Biophys Acta 1829, 1010–4 (2013).

3. Ciro, M. et al. ATAD2 is a novel cofactor for MYC, overexpressed and amplified in aggressive tumors. Cancer Res 69, 8491–8 (2009).

4. Luo, Y. et al. ATAD2 Overexpression Identifies Colorectal Cancer Patients with Poor Prognosis and Drives Proliferation of Cancer Cells. Gastroenterol Res Pract 2015, 936564 (2015).

5. Kalashnikova, E.V. et al. ANCCA/ATAD2 overexpression identifies breast cancer patients with poor prognosis, acting to drive proliferation and survival of triplenegative cells through control of B-Myb and EZH2. Cancer Res 70, 9402–12 (2010).

6. Zheng, L. et al. Oncogene ATAD2 promotes cell proliferation, invasion and migration in cervical cancer. Oncol Rep 33, 2337–44 (2015).

7. Winter-Holt, J.J. et al. Discovery of a Potent and Selective ATAD2 Bromodomain Inhibitor with Antiproliferative Activity in Breast Cancer Models. J Med Chem 65, 3306–3331 (2022).

8. Bamborough, P. et al. A Qualified Success: Discovery of a New Series of ATAD2 Bromodomain Inhibitors with a Novel Binding Mode Using High-Throughput Screening and Hit Qualification. J Med Chem 62, 7506–7525 (2019).

9. Zou, J.X., Revenko, A.S., Li, L.B., Gemo, A.T. & Chen, H.W. ANCCA, an estrogen-regulated AAA+ ATPase coactivator for ERalpha, is required for coregulator occupancy and chromatin modification. Proc Natl Acad Sci U S A 104, 18067–72 (2007).

10. Zou, J.X. et al. Androgen-induced coactivator ANCCA mediates specific androgen receptor signaling in prostate cancer. Cancer Res 69, 3339–46 (2009).

11. Revenko, A.S., Kalashnikova, E.V., Gemo, A.T., Zou, J.X. & Chen, H.W. Chromatin loading of E2F-MLL complex by cancer-associated coregulator ANCCA via reading a specific histone mark. Mol Cell Biol 30, 5260–72 (2010).

12. Cattaneo, M. et al. Lessons from yeast on emerging roles of the ATAD2 protein family in gene regulation and genome organization. Mol Cells 37, 851–6 (2014).

13. Kurat, C.F. et al. Restriction of histone gene transcription to S phase by phosphorylation of a chromatin boundary protein. Genes Dev 25, 2489–501 (2011).

14. Fillingham, J. et al. Two-color cell array screen reveals interdependent roles for histone chaperones and a chromatin boundary regulator in histone gene repression. Mol Cell 35, 340–51 (2009).

15. Gradolatto, A. et al. Saccharomyces cerevisiae Yta7 regulates histone gene expression. Genetics 179, 291–304 (2008).

16. Gal, C. et al. Abo1, a conserved bromodomain AAA-ATPase, maintains global nucleosome occupancy and organisation. EMBO Rep 17, 79–93 (2016).

17. Lombardi, L.M., Ellahi, A. & Rine, J. Direct regulation of nucleosome density by the conserved AAA-ATPase Yta7. Proc Natl Acad Sci U S A 108, E1302–11 (2011).

18. Chacin, E. et al. A CDK-regulated chromatin segregase promoting chromosome replication. Nat Commun 12, 5224 (2021).

19. Cho, C. et al. Structural basis of nucleosome assembly by the Abo1 AAA+ ATPase histone chaperone. Nat Commun 10, 5764 (2019).

20. Zunder, R.M. & Rine, J. Direct interplay among histones, histone chaperones, and a chromatin boundary protein in the control of histone gene expression. Mol Cell Biol 32, 4337–49 (2012).

21. Jambunathan, N. et al. Multiple bromodomain genes are involved in restricting the spread of heterochromatic silencing at the Saccharomyces cerevisiae HMR-tRNA boundary. Genetics 171, 913–22 (2005).

22. Tackett, A.J. et al. Proteomic and genomic characterization of chromatin complexes at a boundary. J Cell Biol 169, 35–47 (2005).

23. Shahnejat-Bushehri, S. & Ehrenhofer-Murray, A.E. The ATAD2/ANCCA homolog Yta7 cooperates with Scm3(HJURP) to deposit Cse4(CENP-A) at the centromere in yeast. Proc Natl Acad Sci U S A 117, 5386–5393 (2020).

24. Morozumi, Y. et al. Atad2 is a generalist facilitator of chromatin dynamics in embryonic stem cells. J Mol Cell Biol 8, 349–62 (2016).

25. Wang, T. et al. ATAD2 controls chromatin-bound HIRA turnover. Life Sci Alliance 4(2021).

26. Koo, S.J. et al. ATAD2 is an epigenetic reader of newly synthesized histone marks during DNA replication. Oncotarget 7, 70323–70335 (2016).

27. Filippakopoulos, P. et al. Histone recognition and large-scale structural analysis of the human bromodomain family. Cell 149, 214–31 (2012).

28. Lloyd, J.T. & Glass, K.C. Biological function and histone recognition of family IV bromodomain-containing proteins. J Cell Physiol 233, 1877–1886 (2018).

29. Khan, Y.A., White, K.I. & Brunger, A.T. The AAA+ superfamily: a review of the structural and mechanistic principles of these molecular machines. Crit Rev Biochem Mol Biol 57, 156–187 (2022).

30. Erzberger, J.P. & Berger, J.M. Evolutionary relationships and structural mechanisms of AAA+ proteins. Annu Rev Biophys Biomol Struct 35, 93–114 (2006).

31. Puchades, C., Sandate, C.R. & Lander, G.C. The molecular principles governing the activity and functional diversity of AAA+ proteins. Nat Rev Mol Cell Biol 21, 43–58 (2020).

32. Glynn, S.E., Kardon, J.R., Mueller-Cajar, O. & Cho, C. AAA+ proteins: converging mechanisms, diverging functions. Nat Struct Mol Biol 27, 515–518 (2020).

33. Xu, Y. et al. Active conformation of the p97-p47 unfoldase complex. Nat Commun 13, 2640 (2022).

34. Gates, S.N. et al. Ratchet-like polypeptide translocation mechanism of the AAA+ disaggregase Hsp104. Science 357, 273–279 (2017).

35. Wang, L., Myasnikov, A., Pan, X. & Walter, P. Structure of the AAA protein Msp1 reveals mechanism of mislocalized membrane protein extraction. Elife 9(2020).

36. Pan, M. et al. Mechanistic insight into substrate processing and allosteric inhibition of human p97. Nat Struct Mol Biol 28, 614–625 (2021).

37. Augustin, S. et al. An intersubunit signaling network coordinates ATP hydrolysis by m-AAA proteases. Mol Cell 35, 574–85 (2009).

38. Zehr, E. et al. Katanin spiral and ring structures shed light on power stroke for microtubule severing. Nat Struct Mol Biol 24, 717–725 (2017).

39. Sandate, C.R., Szyk, A., Zehr, E.A., Lander, G.C. & Roll-Mecak, A. An allosteric network in spastin couples multiple activities required for microtubule severing. Nat Struct Mol Biol 26, 671–678 (2019).

40. Han, H. et al. Structure of spastin bound to a glutamate-rich peptide implies a hand-over-hand mechanism of substrate translocation. J Biol Chem 295, 435–443 (2020).

41. Wang, F., Feng, X., He, Q., Li, H. & Li, H. Structural basis of Yta7 ATPase-mediated nucleosome disassembly. bioRxiv, 2022.05.13.491901 (2022).

42. Koo, S.J. et al. ATAD2 is an epigenetic reader of newly synthesized histone marks during DNA replication. Oncotarget 7, 70323–70335 (2016).

43. Morozumi, Y. et al. Atad2 is a generalist facilitator of chromatin dynamics in embryonic stem cells. J Mol Cell Biol 8, 349–62 (2016).

44. Luger, K., Rechsteiner, T.J. & Richmond, T.J. Expression and purification of recombinant histones and nucleosome reconstitution. Methods Mol Biol 119, 1–16 (1999).

45. Scheres, S.H. RELION: implementation of a Bayesian approach to cryo-EM structure determination. J Struct Biol 180, 519–30 (2012).

46. Punjani, A., Rubinstein, J.L., Fleet, D.J. & Brubaker, M.A. cryoSPARC: algorithms for rapid unsupervised cryo-EM structure determination. Nat Methods 14, 290–296 (2017).

47. Leitner, A., Walzthoeni, T. & Aebersold, R. Lysine-specific chemical cross-linking of protein complexes and identification of cross-linking sites using LC-MS/MS and the xQuest/xProphet software pipeline. Nat Protoc 9, 120–37 (2014).

48. Graham, M., Combe, C., Kolbowski, L. & Rappsilber, J. xiView: A common platform for the downstream analysis of Crosslinking Mass Spectrometry data. bioRxiv, 561829 (2019).

