## Supplementary material for "Structure of the Human ATAD2 AAA+ Histone Chaperone Reveals Mechanism of Regulation and Inter-subunit Communication": Summplementary Information

#### SUPPLEMENTARY FIG 1

a

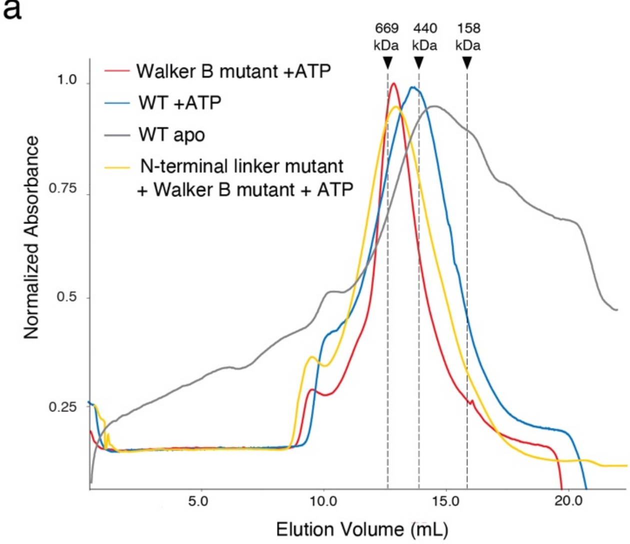


b

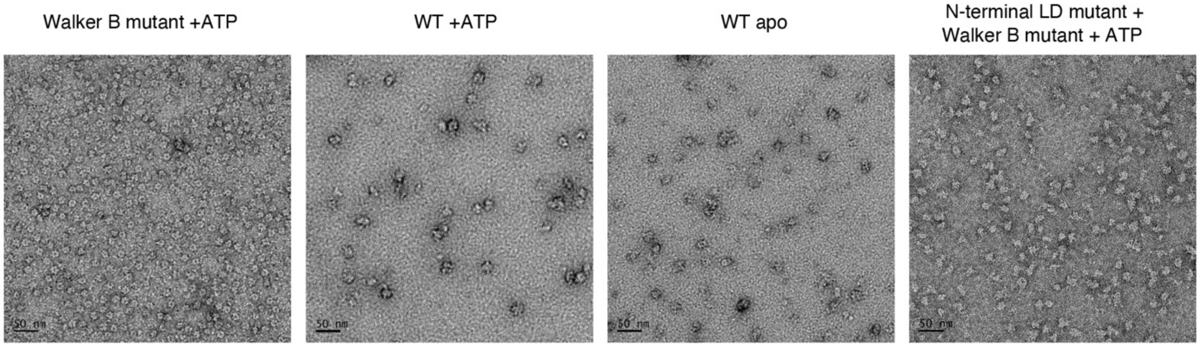


**Supplementary fig 1. Oligomeric assembly of ATAD2**

**a** Gel filtration traces of WT and mutant ATAD2 proteins (Walker B mutant is E532Q and N-terminal LD mutant is D415A/R540A. **b** Negative stain electron micrographs (34,000x magnification) of gel filtration peak fractions of WT and mutant ATAD2 proteins.

#### SUPPLEMENTARY FIG 2

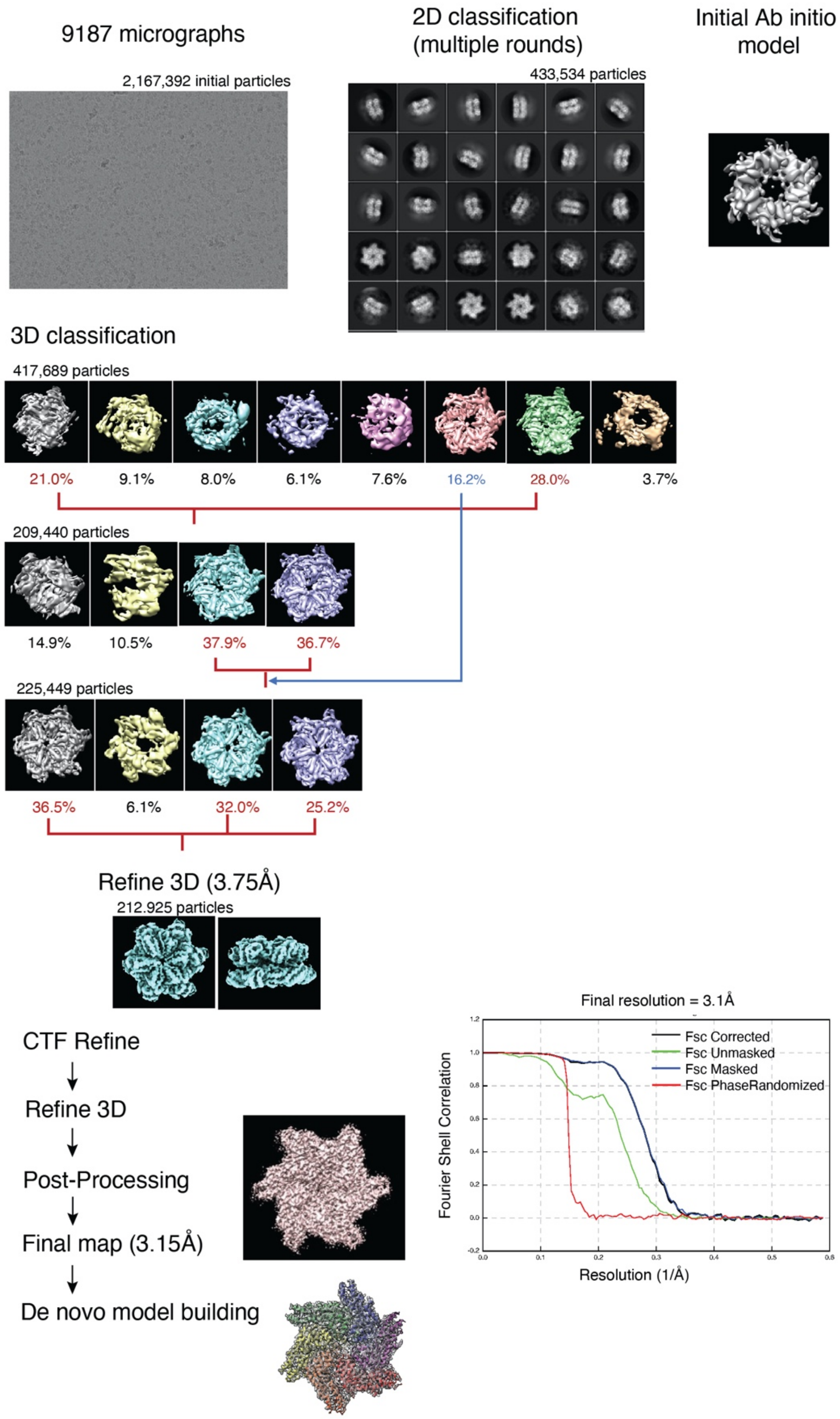

**Supplementary fig 2. Workflow of cryo-EM data processing of ATAD2 Walker B mutant**

Cryo-EM data processing workflow of ATAD2 Walker B mutant + ATP by Relion 3.1, and the Fourier Shell Correlation (FSC) curve of the cryo-EM refinement.

#### SUPPLEMENTARY FIG 3

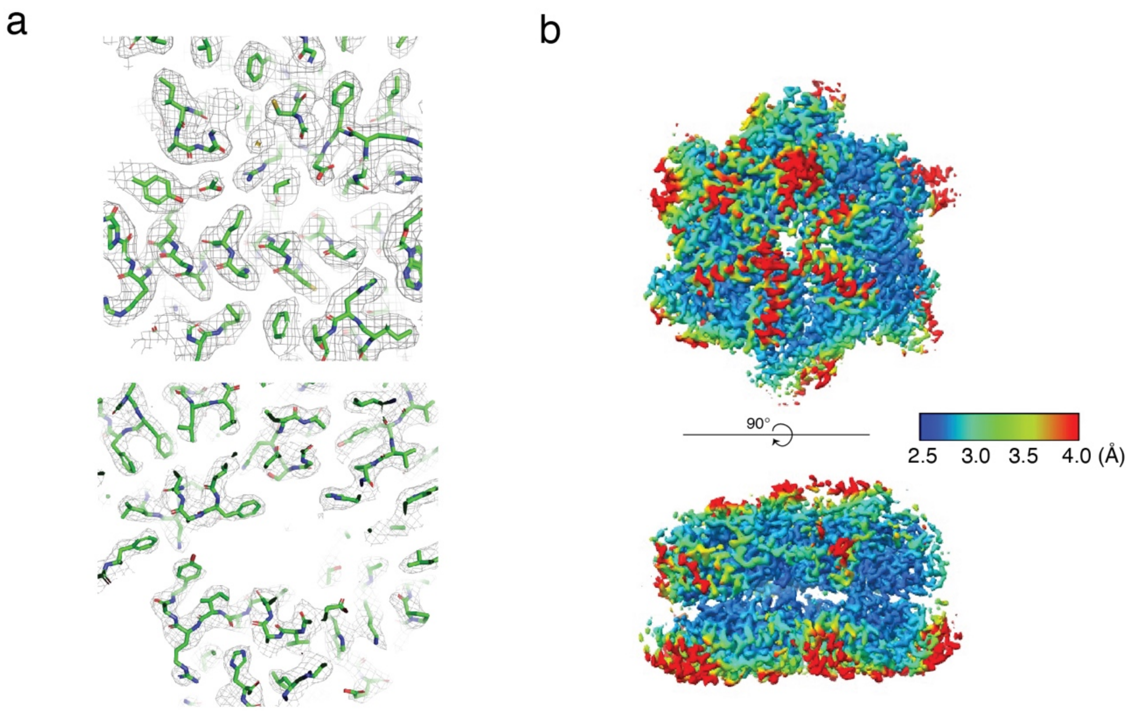


**Supplementary fig 3. Local resolution and model fit of ATAD2 electron density map**

**a** Electron density map of ATAD2 contoured at 3.0 sigma, with model of ATAD2 showing high compatibility with electron density map. **b** Top (top) and side (bottom) views of ATAD2 Walker B mutant cryo-EM maps colored by local resolution. Local resolution map was calculated by final half maps in cryoSPARC.

#### SUPPLEMENTARY FIG 4

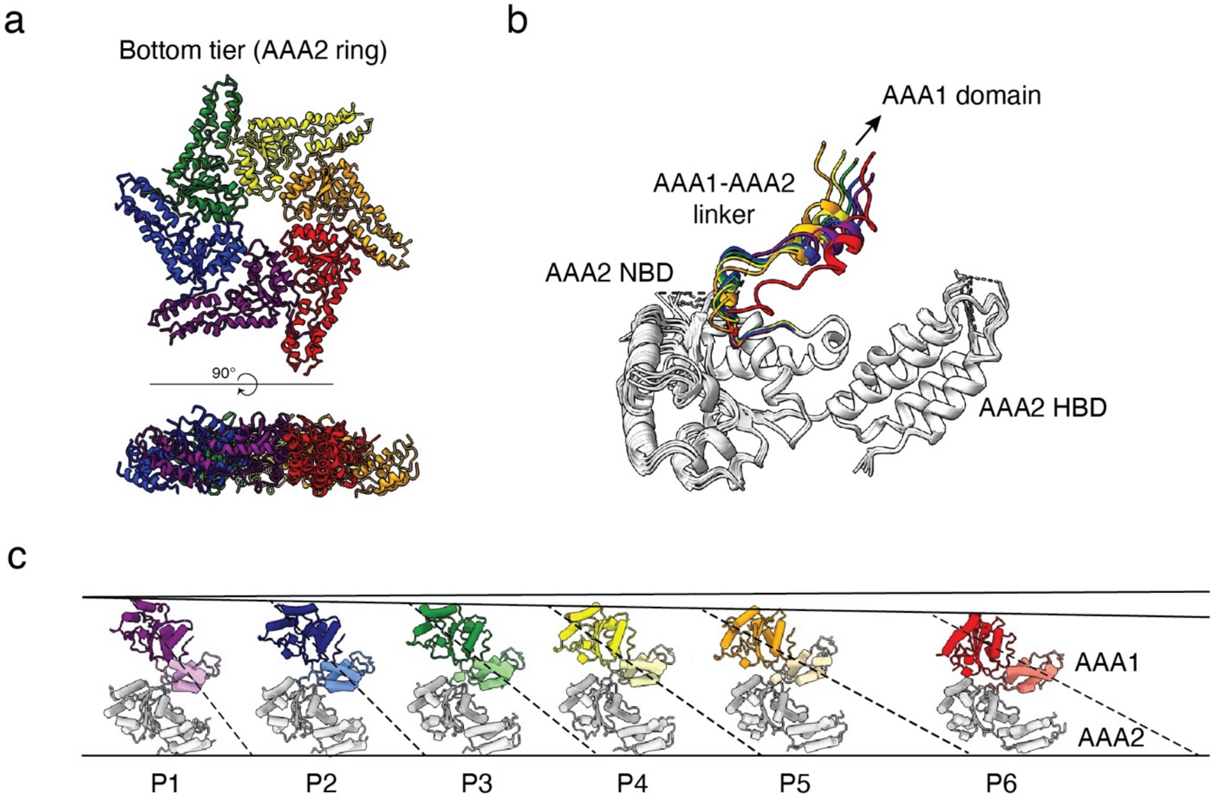


**Supplementary fig 4. Structural elements contributing to ATAD2 hexamer asymmetry**

**a** Symmetric and planar structure of AAA2/C-terminal domain ring. **b** Comparison of AAA1-AAA2 linker conformation in different ATAD2 subunits. Subunits were aligned and superimposed by the AAA2 HBD. Only AAA2 domains and AAA1-2 linkers are shown with AAA1-2 linkers colored according to subunit. **c** Comparison of subunit height and angle of AAA1 (NBD in dark hue, HBD in light hue) with respect to AAA2 (light gray) when subunits are aligned by AAA2 HBD.

#### SUPPLEMENTARY FIG 5

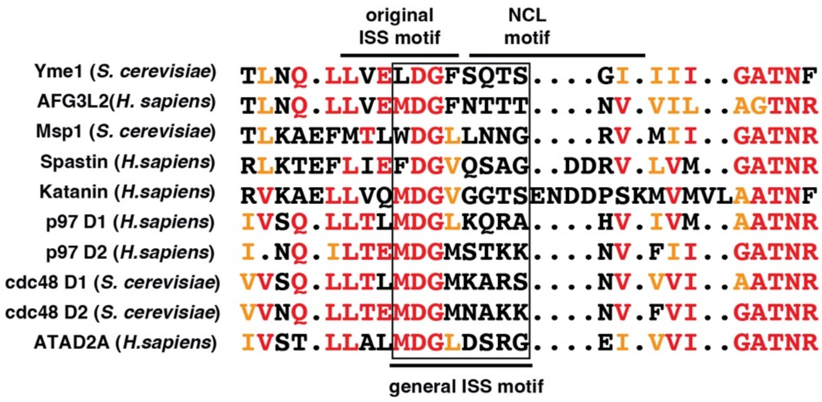


**Supplementary fig 5. Sequence alignment of α3-β4 loop in different AAA+ ATPases**

Alignment of the α3-β4 loop of various AAA+ ATPases, showing the original ISS motif as defined by the DGF tripeptide in the C-terminus of α3 in m-AAA+ proteases (Yme1 and AFG3L2), the NCL motif in meiotic clade AAA+ ATPases (Msp1, Spastin, and Katanin), and an updated "general ISS motif" that applies to the α3-β4 loop of many AAA+ ATPases such as those shown here. Residue positions that are >70% identical are colored red, while positions that are 70%> similar are colored orange.

#### SUPPLEMENTARY FIG 6

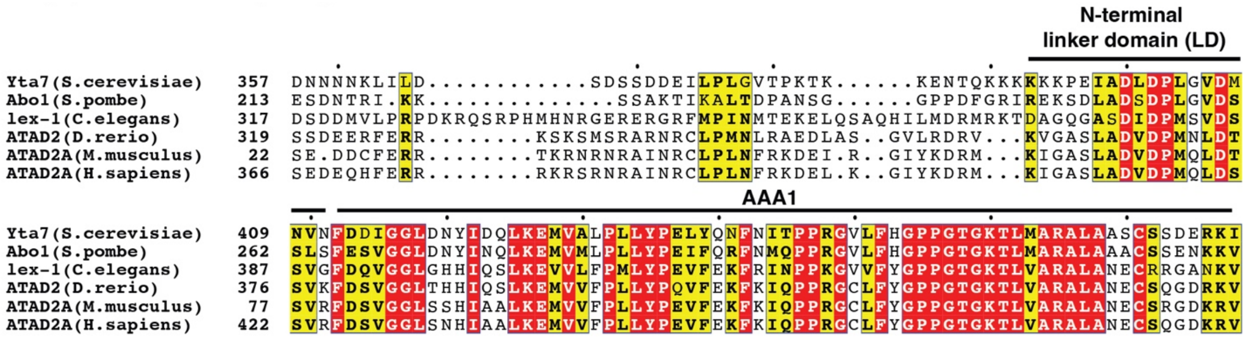


**Supplementary fig 6. Conservation of ATAD2 N-terminal linker domain (LD)**

Multiple sequence alignment of the N-terminal linker domain and AAA1 domains of ATAD2 homologs showing the conservation of the LD. Red represents 100% identity while yellow represents a similarity score of >70%.

#### SUPPLEMENTARY FIG 7

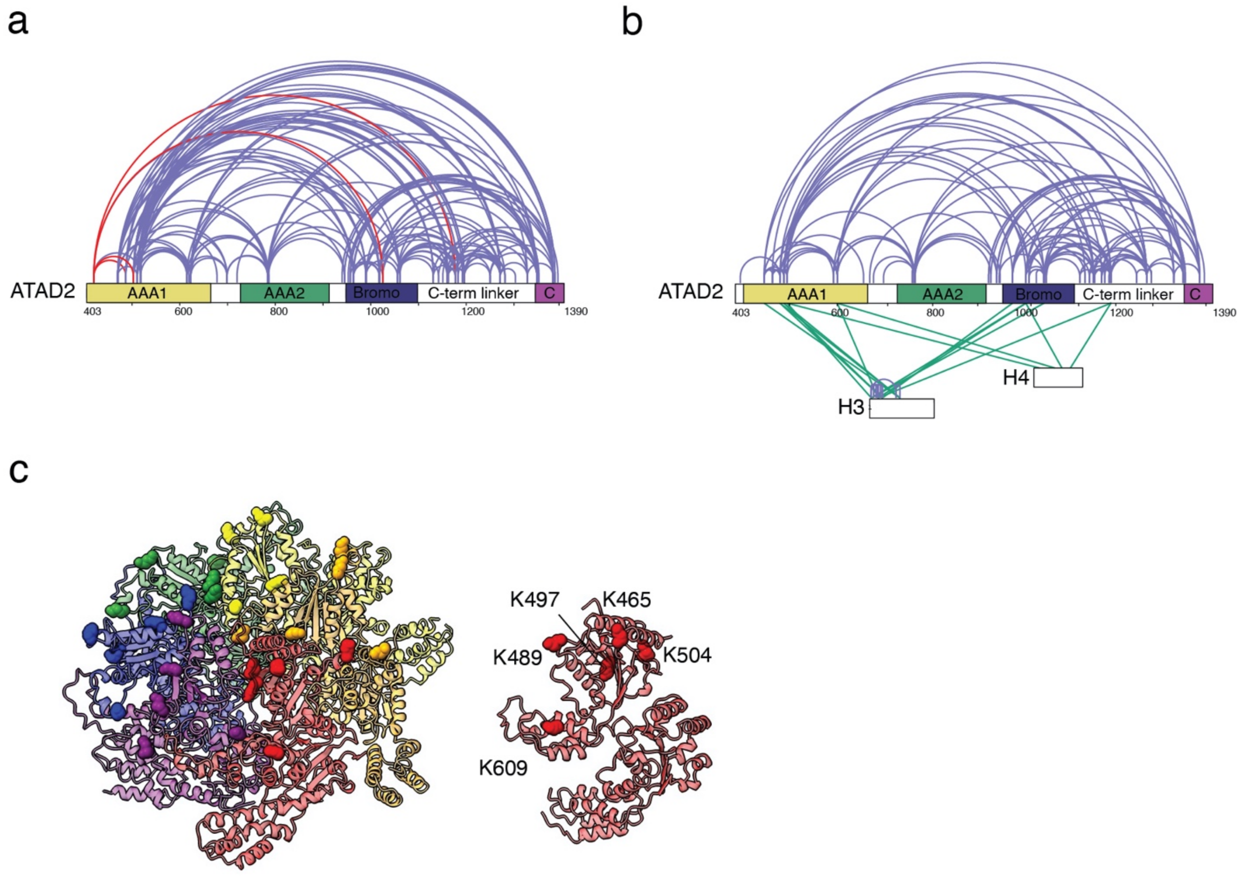


**Supplementary fig 7. Crosslinking Mass Spectrometry of ATAD2-H3H4 complex**

**a** Intramolecular crosslinks of ATAD2 Walker B mutant with ATP. (Crosslinks were analyzed by xQuest/xProphet and filtered by an xQuest LD score cutoff =25 corresponding to an estimated false discover rate < 2%.) Crosslinks of the N-terminal LD that disappear in the ATAD2 Walker B mutant - H3H4 complex are colored red. **b** Intra- and inter-molecular crosslinks of ATAD2 Walker B mutant-H3H4 complex with ATP. **c** ATAD2 residues that crosslink with histone H3H4, mapped onto the ATAD2 hexamer (left), and monomer (right).

### SUPPLEMENTARY FIG 8

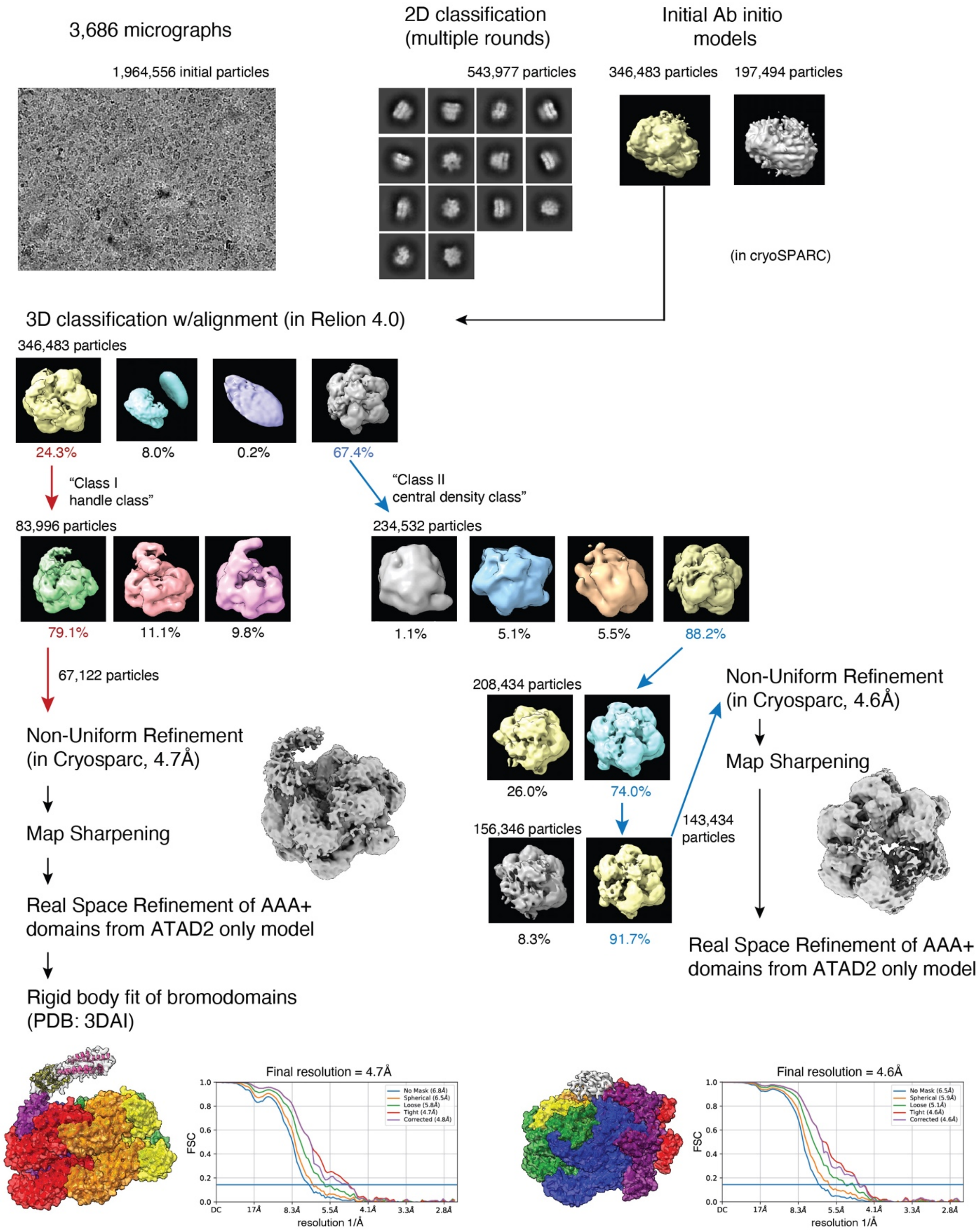

**Supplementary fig 8. Workflow of cryo-EM data processing of ATAD2 Walker B mutant-H3H4 complex**

Cryo-EM data processing workflow of ATAD2 Walker B mutant -H3H4 complex + ATP by cryoSPARC and Relion 4.0, and the Fourier Shell Correlation (FSC) curve of the cryo-EM refinements.

#### SUPPLEMENTARY FIG 9

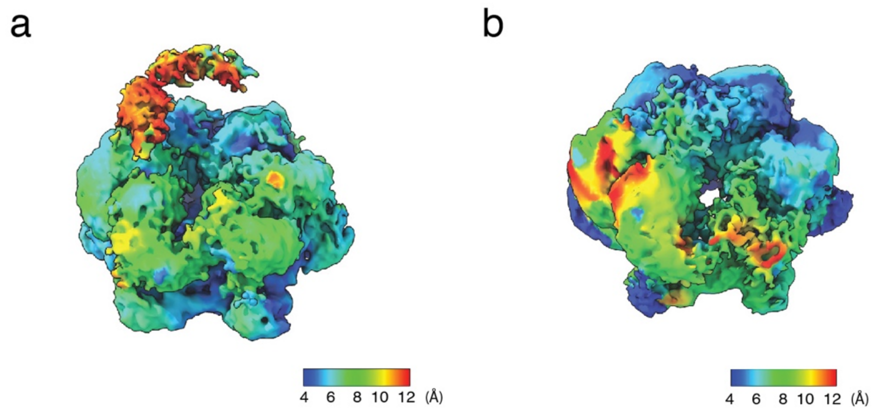


**Supplementary fig 9. Local resolution maps of ATAD2 Walker B mutant-H3H4 complex**

**a,b** Top views of ATAD2 Walker B mutant-H3H4 cryo-EM maps class I (**a**) and class II (**b**) colored by local resolution. Local resolution maps were calculated by final half maps in cryoSPARC.

**Supplementary Movies**

Movie 1 Morph of ATAD2 monomeric subunit structure

Movie 2 Morph of ATAD2 with ATAD2-Histone H3/H4 complex class I structure

Movie 3 Morph of ATAD2 with ATAD2-Histone H3/H4 complex class II structure
